# An Early Miocene skeleton of *Brachydiceratherium* Lavocat, 1951 (Mammalia, Perissodactyla) from the Baikal area, Russia, and a revised phylogeny of Eurasian teleoceratines

**DOI:** 10.1101/2022.07.06.498987

**Authors:** Alexander Sizov, Alexey Klementiev, Pierre-Olivier Antoine

## Abstract

Hippo-like rhinocerotids, or teleoceratines, were a conspicuous component of Holarctic Miocene mammalian faunas, but their phylogenetic relationships remain poorly known. Excavations in lower Miocene deposits of the Olkhon Island (Tagay locality, Eastern Siberia; 16–18 Ma) have opened a unique window on the poorly-known early history of the Lake Baikal ecosystems, notably by unearthing a skeleton of the teleoceratine *Brachydiceratherium shanwangense* (Wang, 1965). The remains provide new insights into the skull and postcranial morphology of this elusive species. The new material is compared with other Eurasian teleoceratines and the relationships within Teleoceratina are investigated through a phylogenetic analysis. *Diaceratherium* Dietrich, 1931 (earliest Miocene, Western Europe) is found to be monotypic and is retrieved as the earliest teleoceratine offshoot. Other genera have more than one species and are also found to be monophyletic, with *Prosantorhinus* Heissig, 1974 (early Miocene, Eurasia) + *Teleoceras* Hatcher, 1894 (Miocene, North America) forming the sister clade of *Brachypotherium* Roger, 1904 (Miocene, Old World) + *Brachydiceratherium* Lavocat, 1951. *Brachydiceratherium* includes eight species spanning the late Oligocene to Late Miocene in Europe and Asia. All teleoceratine genera except *Diaceratherium* span considerable geographical and stratigraphical ranges, likely related to their ultra-generalist ecological preferences.

## INTRODUCTION

Although they are nearly extinct today, rhinocerotids were one of the most widespread and successful groups of large mammals on all the northern continents for over 40 million years. They have occurred across Eurasia and North America since middle Eocene times, and are known from Africa since the Early Miocene (e.g., Prothero et al., 1989; Antoine et al., 2003, accepted; Geraads, 2010). They have also occupied many different locomotory modes, ranging from slender- and long-legged savannah roamers (e.g., elasmotheriines) to hippo-like forms that apparently lived along rivers and lakes (teleoceratines; Prothero et al., 1989; Antoine, 2002). Most hippo-like rhinocerotids are gathered within teleoceratines, a clade at the tribal to sub-tribal level, the phylogenetic relationships of which have never been fully elucidated (Antoine, 2002; Lu et al., 2021). Most teleoceratines had skulls that were either hornless or with a small nasal horn, barrel-shaped bodies, and shortened limb bones. Teleoceratines span the late Oligocene–latest Miocene in Eurasia (Antoine, in press), the Miocene in Africa (Geraads & Miller, 2013), and the early Miocene–early Pliocene in North and Central America (Prothero, 2005). Most of them are interpreted to have been browsers (based on both dental morphology and isotopic studies; MacFadden, 1998; Hullot et al., 2021).

In this study, we describe a skeleton of a teleoceratine from Lower Miocene deposits of Olkhon Island, Lake Baikal area, Siberia. We identify its species assignment and compare it to most teleoceratine species described from Eurasia. This in-depth comparison forms the basis for performing a parsimony analysis of phylogenetic relationships among Eurasian Teleoceratina, and for discussing key events in the paleobiogeography of teleoceratine rhinocerotids.

## LOCALITY AND GEOLOGICAL SETTINGS

Lake Baikal, located in the Baikal Rift System, is morphologically characterised by three basins (Southern, Central and Northern). The Southern and Central basins are thought to have existed permanently since the Paleogene, whereas the Northern Basin did not develop before the Late Miocene (Mats et al., 2010, 2011). Olkhon Island (Russian: Ольхон) is located in the transitional zone between the Central and the Northern basins of Lake Baikal. It is separated from the mainland in the west by a shallow Maloe More strait (Russian: Малое Море; in English literally the Small Sea) of the Northern Basin that extends far to the south. In the south, Maloe More strait is connected through the narrow Olkhonskie Vorota strait (Russian: Ольхонские Ворота; in English literally the Olkhon Gate) to the central part of Lake Baikal. From the northwestern part of Olkhon Island, one locality known as Tagay or Tagai (Russian: Тагай or Тогай) has yielded numerous terrestrial fossils of the Neogene (Fig. 1). The Neogene sediments in Tagay Bay belong to the Tagay Formation (Logachev et al., 1964; Mats et al., 2001; Mats, 2013, 2015). Sediments are exposed in the northeastern part of the bay in a steep erosional cliff up to 15 m high. Elsewhere along the shores of the bay, this cliff is levelled by landslides. The cliff borders a large landslide cirque and a sandy beach below.

**Fig. 1.**
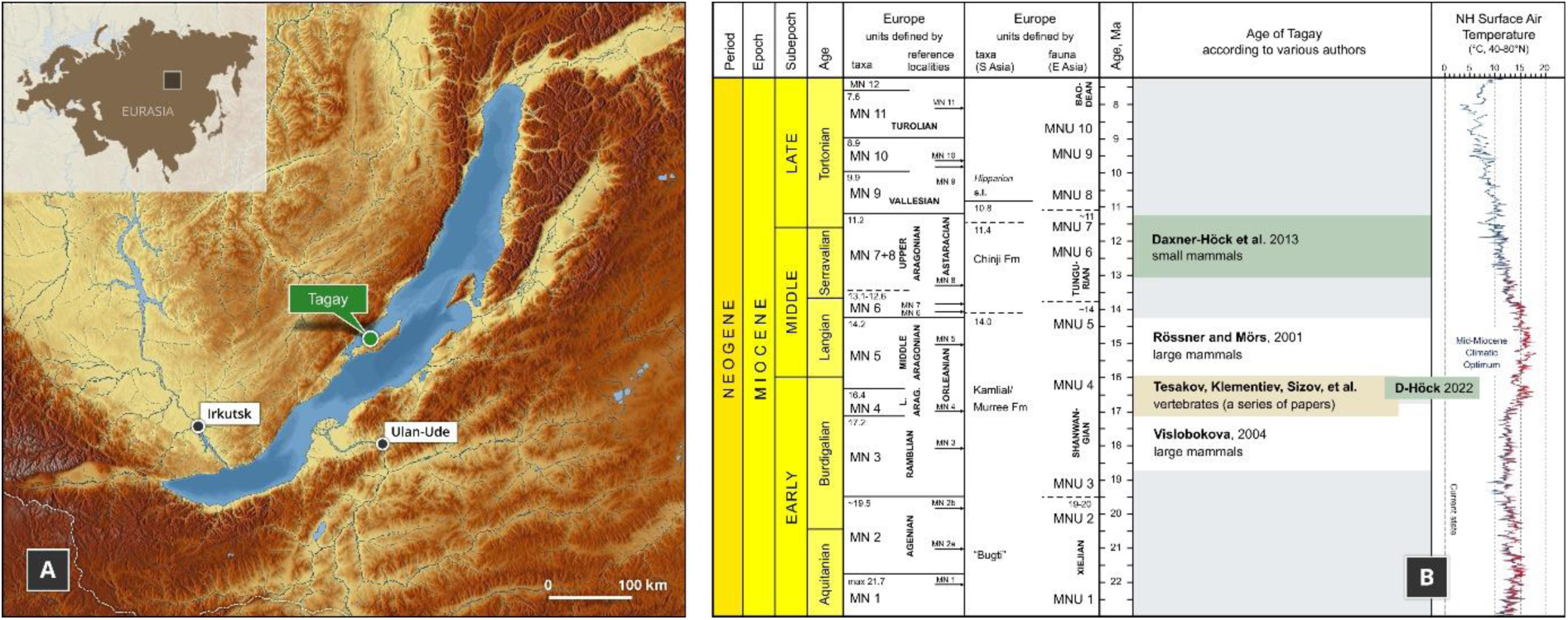
Geographic position of Tagay locality on Olkhon Island, Baikal Region, Russia (A) and age of Tagay Formation according to various authors (B).

The Tagay locality was discovered in the 1950s (Kitainik & Ivaniev, 1958). The first paleontological studies of the large mammals were performed in 1958 under the direction of N.A. Logachev (Logachev et al., 1964). Studies of small mammals have been carried out since the 1970s (Pokatilov, 2004; Daxner-Höck et al., 2022). Tagay preserves an abundant fossil fauna that includes molluscs and vertebrates such as fish, amphibians, reptiles, birds and mammals. However, a significant part of the paleontological material was determined for the longest time only tentatively: Mustelidae and Felidae among carnivorans, *Anchitherium* sp, *Metaschizotherium*(?) sp., and *Dicerorhinus*(?) sp. among perissodactyls, *Palaeomeryx* sp. and Bovidae indet. among artiodactyls (Logachev et al., 1964). The reexamination of artiodactyl remains led to the identification of Cervidae (*Amphitragulus boulangeri*, *Lagomeryx parvulus*, *Stephanocemas* sp.), Palaeomerycidae (*Orygotherium tagaiense*, *Palaeomeryx* cf. *kaupi*) and Anthracotheriidae (*Brachyodus intermedius*) (Vislobokova, 1990, 1994, 2004). Chelonians were studied by Khosatzky and Chkhikvadze (1993) and the ichthyofauna by Filippov & Sytchevskaya (2000).

Small mammals from the Tagay-1 section were recently revised, with a list of 21 taxa documenting erinaceids, talpids, plesiosoricids, and soricids among eulipotyphlans, palaeolagid lagomorphs, and sciurids, aplodontids, mylagaulids, glirids, castorids, eomyids, and cricetodontine muroids among rodents (Daxner-Höck et al., 2022).

The sedimentological, stratigraphical, and palaeontological aspects of the sediments were described by Kossler (2003). A new phase of the study of Tagay locality started in 2008, with numerous publications since that time (Rage & Danilov, 2008; Klementiev, 2009; Danilov et al., 2012; Syromyatnikova, 2014, 2015; Tesakov & Lopatin, 2015; Klementiev & Sizov, 2015; Zelenkov, 2016; Sotnikova et al, 2021).

The Tagay Formation consists of alternating beds of clays and clayey sands containing interlayers and lenses of carbonate concretions of diagenetic origin. Deposits rest upon the crystalline basement, and are submerged below water to the south. Clay beds are mostly green and brown, sometimes black. There are also lenses and interlayers of brick red and red clay and loam. Bone beds, deposited in successive sedimentary cycles, were given letters from top to bottom (i.e., downsection: A–H; Fig. 2 A, B). Most clay beds have predominant ferruginous-magnesian montmorillonites composition. A remarkable feature of these clay sediments is the high (up to 8%) content of silt-psammite-psephite admixtures. Moreover, most psammite-psephitic fragments are not rounded and have angular and indented outlines, which indicates a lack of transportation. The lithology of the sections and further details on the bone beds were described in other studies (Logachev et al., 1964; Sizov & Klementiev, 2015).

**Fig. 2.**
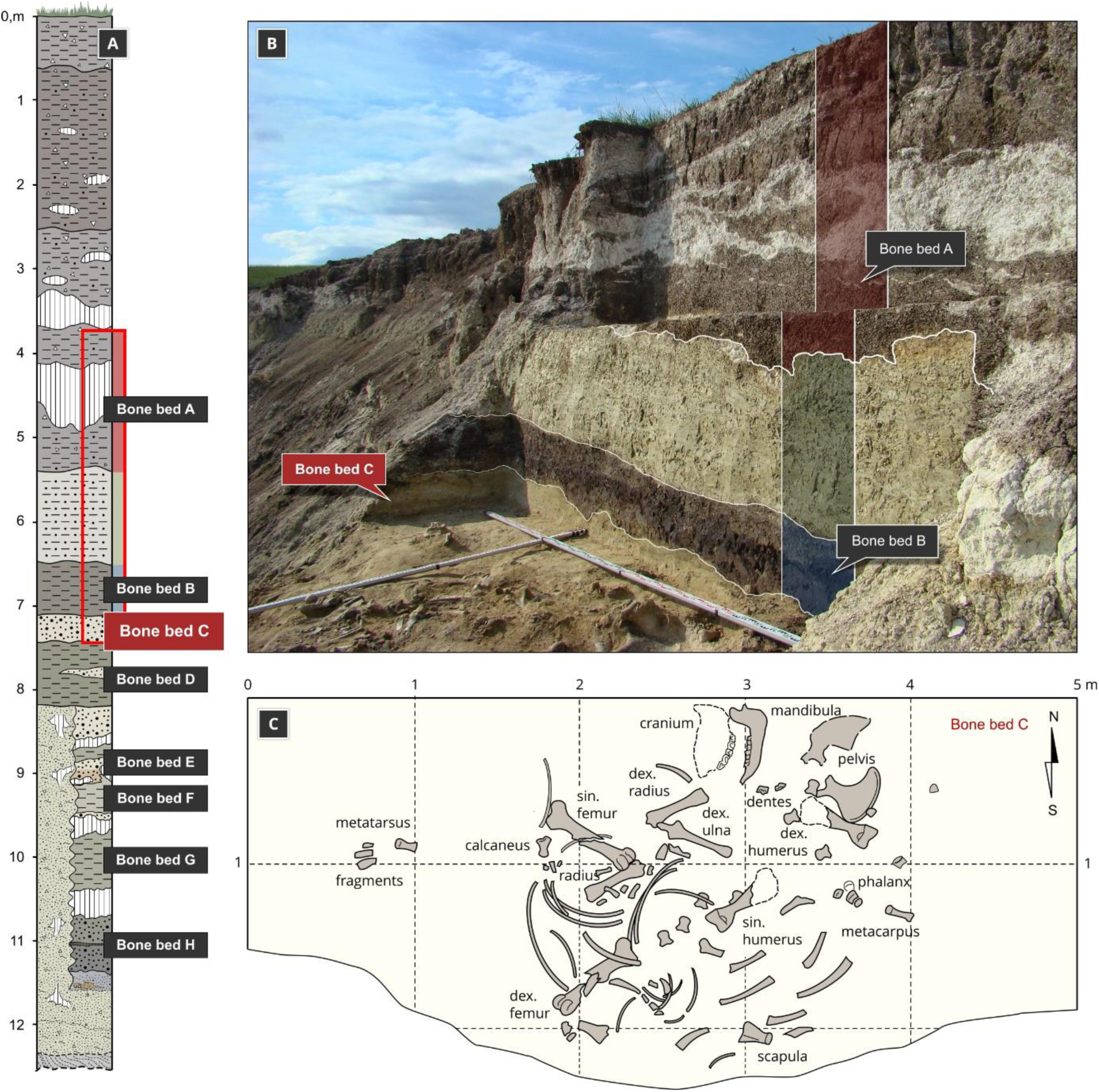
Geological structure of the Tagay section (A), photo (B) and plan (C) of the excavations of the Miocene rhinocerotid at Tagay site in 2008 (Olkhon Island, Baikal Region, Russia).

Neogene continental deposits in the late early Miocene Tagay locality have yielded a diverse vertebrate fauna. Following most works, the age of the Tagay Fauna correlates to the European Mammal Zones MN3 through MN5 (20–15 Ma; Rössner & Mörs, 2001; Vislobokova, 2004; Klementiev & Sizov, 2015; Sotnikova et al., 2021). Other researchers have correlated the Tagay fauna or to MN 7+8 and Chinese Mammal Unit NMU7 (13–11 Ma; Daxner-Höck et al., 2013). More recently, Daxner-Höck et al. (2022) proposed a more precise age of ∼16.5-16.3 Ma based on micromammalian bicostratigraphy and the magnetic polarity pattern of the Tagay-1 section (Fig. 1 B), which is in agreement with our preferred interval.

## MATERIAL AND METHODS

All the remains described here belong to a single adult individual (IZK79-1-08C-1/), stored in the collection of the Institute of the Earth’s Crust (Irkutsk, Russia). Alexey Klementiev and Gennady Turkin discovered this skeleton in 2008 at Tagay (Fig. 2 B, C) (Klementiev, 2009). Capital letters are used for upper teeth (I, C, D, P, M), and lower-case letters for lower teeth (i, c, d, p, m). Rhinocerotid dental terminology follows Heissig (1969, 1972a: pl. 13) and Antoine (2002), while dental and skeletal measurements were taken according to Guérin (1980). Anatomical descriptions follow basically the same sequence as in Antoine (2002), and Antoine et al. (2010). Dimensions are given in mm.

The stratigraphical framework is based on the Neogene geological time scale and European Land Mammal Ages (Hilgen, Lourense & Van Dam, 2012; Raffi et al, 2020).

### 3D-rendering

All bones of the rhinoceros were scanned with a resolution of 0.25 mm using a RangeVision Smart 3D scanner. RangeVision Smart has three areas of scanning and is equipped with colour cameras 1.3 megapixels. We used the associated RangeVision 2020.2 software for visualization, segmentation and 3D rendering.

### Phylogenetic analysis

The phylogenetic analysis was based on 282 cranio-mandibular, dental, and postcranial characters primarily derived from the dataset of Antoine (2002, 2003) which was scored on 31 ceratomorph species (one tapirid and 30 rhinocerotoids). All multistate characters were treated as additive, except for the characters 72, 94, 102, 140, 187, and 269 (non-additive; as in Antoine, 2002).

The living Brazilian tapir *Tapirus terrestris* (Linnaeus, 1758), the Eocene non-rhinocerotid rhinocerotoid *Hyrachyus eximius* Leidy, 1871 and the Paleogene stem rhinocerotids *Trigonias osborni* Lucas, 1900 (Eocene of North America) and *Ronzotherium filholi* (Osborn, 1900) (Oligocene of Western Europe) were selected as outgroups. We also included a branching group (Antoine, 2002, 2003; Orliac et al., 2010; Boivin et al., 2019), consisting of non-teleoceratine taxa, with the aim of testing the monophyly and relationships of the Teleoceratina among the Rhinocerotinae. These consist of 12 species, including an early-diverging representative of Rhinocerotinae (*Plesiaceratherium mirallesi* (Crusafont, Villalta & Truyols, 1955)), three species of Aceratheriini (*Aceratherium incisivum* Kaup, 1832, *Acerorhinus zernowi* (Borissiak, 1914), and *Alicornops simorrense* (Lartet, 1851)), and eight members of the Rhinocerotina, encompassing all five living rhinoceroses, namely the Indian rhino (*Rhinoceros unicornis* Linnaeus, 1758), the Javan rhino (*Rhinoceros sondaicus* Desmarest, 1822), the Sumatran rhino (*Dicerorhinus sumatrensis* (Fischer, 1814)), the white rhino (*Ceratotherium simum* (Burchell, 1817)), and the black rhino (*Diceros bicornis* (Linnaeus, 1758)), in addition to three fossil species: *Lartetotherium sansaniense* (Lartet in Laurillard, 1848) (Miocene of Europe; Heissig, 2012), *Gaindatherium browni* Colbert, 1934 (Miocene of South Asia; Heissig, 1972a; Antoine, in press), and *Nesorhinu*s *philippinensis* (Von Koenigswald, 1956) (early Middle Pleistocene of the Philippines; Antoine et al., 2022 and references therein).

The ingroup sensu stricto (Teleoceratina) comprises 15 taxa, with *Teleoceras fossiger* Cope, 1878 (late Miocene to earliest Pliocene, North America), *Brachypotherium brachypus* (Lartet in Laurillard, 1848) (late early and middle Miocene, Eurasia), *Brachypotherium perimense* (Falconer & Cautley, 1847) (Miocene, South Asia), *Prosantorhinus germanicus* (Wang, 1929) (late early and middle Miocene, Europe), *Prosantorhinus douvillei* (Osborn, 1900) (late early and early middle Miocene, Europe), *Prosantorhinus laubei* Heissig & Fejfar, 2007 (early Miocene, central Europe), and a comprehensive sample of taxa either classically or more recently assigned to *Diaceratherium* Dietrich, 1931. These consist of the type species *D*. *tomerdingense* Dietrich, 1931 from the earliest Miocene of Tomerdingen (Germany), *D*. *lemanense* (Pomel, 1853) from the latest Oligocene-early Miocene of Western Europe (also described under the *Diceratherium* (*Brachydiceratherium*) *lemanense* combination by Lavocat, 1951), *D*. *aurelianense* (Nouel, 1866) from the early Miocene of Western Europe. The taxonomic sample also includes *D*. *asphaltense* (Depéret & Douxami, 1902) from the earliest Miocene of Western Europe, *D*. *fatehjangense* (Pilgrim, 1910), from the Miocene of Pakistan and early Miocene of Kazakhstan (previously described as “*Brachypotherium aurelianense* Nouel, var. nov. *Gailiti*” by Borissiak, 1927), and *D*. *aginense* (Répelin, 1917) from the earliest Miocene of Western Europe. Lastly, we have considered *D*. *shanwangense* (Wang, 1965) from the late early Miocene of eastern China (Shanwang; Lu et al., 2021), Japan, and eastern Siberia (Tagay; this work), and *D*. *lamilloquense* Michel, in Brunet et al., 1987 from the late Oligocene of France. We also included *Aceratherium gajense intermedium* Lydekker, 1884, which has disputed taxonomic affinities. It was previously assigned to the aceratheriine genera *Subchilotherium* (e.g., Heissig, 1972a) or *Chilotherium* (e.g., Khan et al., 2011), although based on a parsimony analysis taking into account the holotype and original hypodigm, Antoine et al. (2003) considered that it might be a teleoceratine instead, of uncertain generic assignment. The recognition of associated dental and postcranial remains from the Potwar Plateau (late early to early late Miocene, Pakistan) allowed for defining the new combination *Diaceratherium intermedium* (Lydekker, 1884) for this taxon, as recently proposed by Antoine (in press).

Three representatives of Teleoceratina, *Diaceratherium* cf. *lamilloquense* from the late Oligocene of Thailand (Marivaux et al., 2004), *Brachypotherium gajense* (Pilgrim, 1910), from the late Oligocene–earliest Miocene of Pakistan, and *Prosantorhinus shahbazi* (Pilgrim, 1910), from the early Miocene of Pakistan (combinations proposed by Antoine et al., 2010 and Antoine, in press) were not included in the analysis, due to their poorly-known hypodigms, restricted to a few elements.

Moreover, *Diaceratherium askazansorense* Kordikova, 2001 from the early Miocene of Kazakhstan was not included, as dental and postcranial elements assigned to this taxon closely resemble those of *Pleuroceros blanfordi*, a stem member of Rhinocerotinae (early Miocene of South Asia; Antoine et al., 2010; Prieto et al., 2018) and, to a lesser extent, of *Pleuroceros pleuroceros* (earliest Miocene of western Europe; Antoine et al., 2010; Antoine & Becker, 2013). Similarly, the early late Oligocene species *Diaceratherium massiliae* Ménouret & Guérin, 2009 was recently shown to be a junior synonym of the short-limbed and early-diverging rhinocerotid *Ronzotherium romani* Kretzoi, 1940, through a re-examination of most available material and the recognition of new associated dental and postcranial specimens in Switzerland (Tissier et al., 2021).

Details on the specimens, collections, direct observation and/or literature used for scoring taxa, with references used) are provided as taxon notes in the morphological data matrix (supplementary file S3). The parsimony analyses were performed through the heuristic search of PAUP 4 3.99.169.0 (Swofford, 2002), with tree-bisection-reconnection (reconnection limit = 8), 1000 replications with random addition sequence (10 trees held at each step), gaps treated as missing, and no differential weighting or topological constraints. Branch support was estimated through Bremer indices (Bremer, 1994), also calculated in PAUP 4 3.99.169.0.

### Systematics

Generic and suprageneric systematics follow the present parsimony analysis (see below).

## SYSTEMATIC PALAEONTOLOGY

Order Perissodactyla OWEN, 1848

Family Rhinocerotidae GRAY, 1821

Subfamily Rhinocerotinae GRAY, 1821

Tribe Rhinocerotini GRAY, 1821

Subtribe Teleoceratina HAY, 1902

Genus *Brachydiceratherium* LAVOCAT, 1951

Syn. *Diaceratherium* DIETRICH, 1931 (partim)

### Type species

*Acerotherium lemanense* Pomel, 1853 by subsequent designation (Lavocat, 1951)

### Included species

*Rhinoceros aurelianensis* Nouel, 1866 from the early Miocene of Western Europe; *Aceratherium intermedium* Lydekker, 1884, from the early–late Miocene of the Indian Subcontinent and China (Deng and Gao, 2006; Antoine et al., 2013; Antoine, in press); *Diceratherium asphaltense* Depéret & Douxami, 1902 from the earliest Miocene of Western Europe; *Teleoceras fatehjangense* Pilgrim, 1910, from the Miocene of Pakistan and early Miocene of Kazakhstan (senior synonym of “*Brachypotherium aurelianense* Nouel, var. nov. *Gailiti*” by Borissiak, 1927); *Teleoceras aginense* Répelin, 1917 from the earliest Miocene of Western Europe; *Plesiaceratherium shanwangense* Wang, 1965 from the late early Miocene of eastern China (Shanwang; Lu et al., 2021), Japan, and eastern Siberia (Tagay; this work); *Diaceratherium lamilloquense* Michel, in Brunet et al., 1987 from the late Oligocene of France.

### Diagnosis

Teleoceratines with a small nuchal tubercle, articular tubercle smooth on the squamosal, with cement present on cheek teeth, protocone always constricted on P3-4, labial cingulum usually absent on lower premolars and always present on lower molars, foramen vertebrale lateralis present and axis-facets transversally concave on the atlas, a postero-distal apophysis low on the tibia, and a latero-distal gutter located posteriorly on the fibula.

Distinct from *Diaceratherium tomerdingense* in possessing a long metaloph on M1-2, no mesostyle on M2, a distal gutter on the humeral epicondyle, an anterior side of the semilunate with a sharp distal border, no posterior expansion on the pyramidal-facet of the unciform, and a trapezium-facet present on the McII.

Differs from representatives of *Brachypotherium* in having close parietal crests, a median ridge on the occipital condyles, a mandibular symphysis less massive, a labial cingulum usually or always absent on upper premolars, an external groove developed on the ectolophid of lower cheek teeth, a V-shaped lingual opening of the posterior valley of lower premolars (in lingual view), a paraconid developed on p2, no second distal radius-ulna facet, a symmetric semilunate-pyramidal distal facet, a posterior McIII-facet present on the McII, and a fibula-facet subvertical on the astragalus.

Distinct from species referred to as *Prosantorhinus* in showing no latero-ventral apophysis on the nasals, close fronto-parietal crests, ad no posterior groove on the processus zygomaticus of the squamosal.

### Geographical and stratigraphical range

Late Oligocene and Miocene of Eurasia, with an Early Miocene climax. *Brachydiceratherium shanwangense* (Wang, 1965) See synonymy list in Lu et al. (2021)

### Holotype

IVPP V 3026, left maxilla with upper cheek tooth series (P2-M3), stored at the Institute of Vertebrate Paleontology and Paleoanthropology, Chinese Academy of Sciences (IVPP).

### Stratum typicum and locus typicus

Early Miocene (Shanwang Formation, Shanwangian Age/Stage of China, middle Burdigalian); Xiejiahe locality, Shanwang Basin, Shandong, China (see Lu et al., 2021).

### Diagnosis

Representative of *Brachydiceratherium* with a lateral apophysis present on the nasals, a median nasal horn present on the nasals, premolar series short with respect to the molar series, roots distinct on the cheek teeth, crochet always simple and lingual cingulum usually absent and always reduced on P2-4, crista always present on P3, protocone strongly constricted on M1-2, lingual cingulum usually absent on lower premolars and always absent on lower molars, d1/p1 absent in adults, glenoid fossa with a medial border straight on the scapula, distal gutter absent on the lateral epicondyle of the humerus, proximal radius-ulna facets always fused, and trochanter major low on the femur.

Distinguished from *Bd. lamilloquense*, *Bd. lemanense*, *Bd. asphaltense*, and/or *Bd. aurelianense* in having I1s oval in cross section, no labial cingulum on upper cheek teeth, a strong paracone fold on M1-2 and a constricted hypocone on M1, M3s with a triangular occlusal outline, a radius with a high posterior expansion of the scaphoid-facet, a femoral head hemispheric, an astragalus with a laterodistal expansion, very low-and-smooth intermediate reliefs on metapodials, and a long insertion of m. interossei on lateral metapodials.

Differs from *Bd. aginense* in having a processus postorbitalis on the frontal bone and a median ridge on the occipital condyle, but no posterior groove on the processus zygomaticus of the squamosal, molariform P2s (protocone and hypocone lingually separate), a long metaloph on M1-2, a posterior groove on M3, a shallow gutter for the m. extensor carpi on the radius, a posterior MtII-MtIII facet developed, but no cuboid-MtIII contact.

Distinct from *D. intermedium* in showing usually a lingual cingulum on upper molars, a strong paracone fold on M1-2, a lingual cingulum usually absent on lower premolars, and a right angle between the cuboid-facet and the base of the tuber calcanei on the calcaneus Differs from *Bd. lemanense* in possessing a low zygomatic arch with a processus postorbitalis, a small processus posttympanicus and a well-developed processus paraoccipitalis. Distinct from *Bd. asphaltense* in having closer fronto-parietal crests and a brachycephalic shape.

Differs from *D. lamilloquense* in showing a protoloph joined to the ectoloph on P2 and molariform P3-4s (protocone and hypocone lingually separate). Differs from *D. aurelianense* in having no metaloph constriction on P2-4 and a protocone weakly developed on P2.

### Geographical and stratigraphical range

Late early Miocene of the Shanwang Basin, Shandong Province, China (see Lu et al., 2021) and of Irkutsk Region, Russia (Tagay locality, Olkhon Island, Lake Baikal).

### Material studied

IZK79-1-08C-1, almost complete skeleton, including the skull (occipital, parietal, frontal, the right zygomatic and lacrimal, both nasals, and temporals with processes and also premaxillae), the jaws, most vertebrae and ribs, both humeri, radii and ulnae, both femora, tibiae, right fibula, most metacarpals, and several metatarsals and phalanges. The skeleton described herein was found disarticulated at the junction of layers of sand and clay (Fig. 2 B, C). In general, the right side of the individual is much better preserved than the left one.

## DESCRIPTION

### Skull

The skull (Fig. 3, Table 1) was found disarticulated, but there is no doubt that the separate bones belong to the same individual, because they were found in close proximity to one another with no extraneous elements, and they fit together well. The temporal, zygomatic and lacrimal, nasal, frontal, parietal and occipital fit each other perfectly. The remaining bones are matching in size, colour and texture. The skull is short and relatively wide (Length from condyles to nasals = 540 mm, Width at the frontals ≈ 190 mm), belonging to a large-sized adult rhinocerotid. The separated nasal bones are long and longer than the preserved part of the premaxilla, relatively thin and bear a lateral apophysis. Roughness for a small nasal horn is preserved at the tip of the nasals. In lateral view, the foramen infraorbitalis and the posterior border of the U-shaped nasal notch are both located above the P3, while the anterior border of the orbit is above the M1. The minimum distance between the posterior edge of the nasal notch and the anterior border of the orbit is 67.2 mm.

**Fig. 3.**
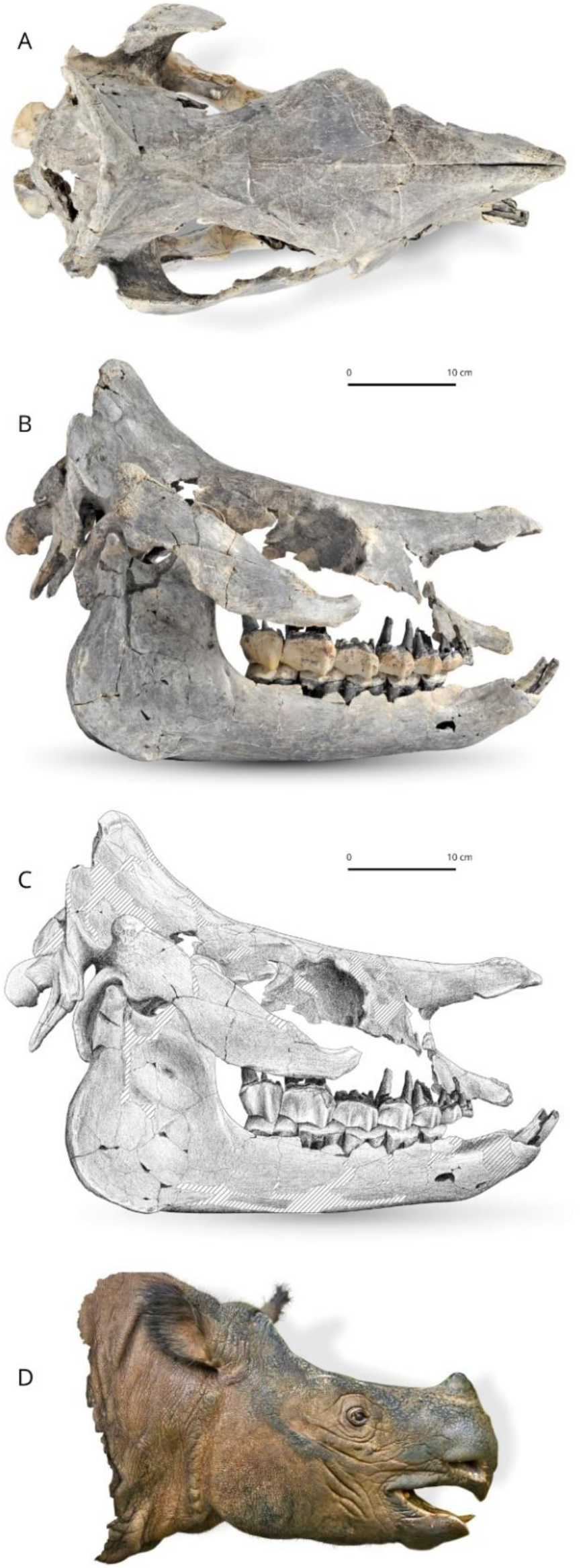
*Brachydiceratherium shanwangense* (Wang, 1965) from Tagay, Baikal Region, Russia, late early Miocene. Photos in dorsal (A) and lateral views (B) of the skull and mandible IZK79-1-08C-1. C - Scientific drawing of the right lateral view of the skull (based on B). Striped areas are reconstructed. D – Tentative reconstruction of the head in lateral view, by one of us (AS).

**Table 1.** Cranial measurements of *Brachydiceratherium shanwangense*, from Tagay, early Miocene of Eastern Siberia, in mm. 1, Length (occipital-premaxilla distance); 2, Length (occipital-nasal distance); 3, Upper length (nasal-occipital crest distance); 4, Nasal incisure length; 5, Minimal width; 6, Occipital crest-postorbital process distance; 8, Occipital crest-lacrimal process distance; 9, Nasal incisure-orbit distance; 13, Post-M3-condyle distance; 14, Nasal-orbit distance; 15, Occipital crest width; 16, Mastoid apophyses width; 17, Inter frontoparietal crest minimal distance; 18, Postorbital process width; 19, Lacrimal process width; 21, Zygomatic width; 22, Nasal incisure width; 23, Occipital height; 25, P2-level height; 26, P4-M1-level height; 27, M3-level height; 31, Foramen magnum width; 32, Inter-occipital condyle width. Numbers coincide with measurements as defined and illustrated by Guérin (1980, fig. 1, table 1).

| 1 | 2 | 3 | 4 | 5 | 6 | 7 | 8 | 9 | 10 | 11 | 12 | 13 | 14 | 15 | 16 |
| --- | --- | --- | --- | --- | --- | --- | --- | --- | --- | --- | --- | --- | --- | --- | --- |
| 505 | 540 | 455.8 | 174 | 125 | 249.6 |  | 293.6 | 72.6 |  |  |  | 254 | 219 | 160 | 224.8 |

| 17 | 18 | 19 | 20 | 21 | 22 | 23 | 24 | 25 | 26 | 27 | 28 | 29 | 30 | 31 | 32 |
| --- | --- | --- | --- | --- | --- | --- | --- | --- | --- | --- | --- | --- | --- | --- | --- |
| 26.7 | 179 | 189 |  | 308.2 | 69.7 | 144.6 |  | 140 | 155 | 156 |  |  |  | 43 | 112.4 |

### Cranial features

The skull was partly destroyed and some elements were reconstructed in anatomical position by one of us (AS). It is short, broad, and elevated. The dorsal profile of the skull is concave, with a small protuberance for a short nasal horn and an upraised parietal bone (50°). In lateral view, the nasals have a small ventrolateral prominence (lateral apophysis, sensu Antoine, 2002). The maxilla is badly damaged and the area of the foramen infraorbitalis is restored on both sides. Nevertheless, based on the preserved part of the maxilla, a position above P4 can be hypothesised. The posterior end of the nasal notch is located above the anterior part of P3. The nasal septum is not ossified at all. The premaxillae are broken rostrally. They form a short and elevated strip, slightly dipping frontward, with a deep ventral sulcus. Relations between nasal and lacrimal bones are not observable, and neither are the lacrimal processi. The anterior border of the orbit is situated above the middle of M1. On the frontal, a pair of smooth tubercles lay on the dorsal and posterodorsal edges of the orbit (processus postorbitalis). The anterior base of the processus zygomaticus maxillari is low, ∼1 cm above the neckline of molars. The zygomatic arch forms a straight, low, and oblique strip, with parallel dorsal and ventral borders. It is parallel to the dorsal outline of the skull, with a rounded and rugose posterodorsal tip. A marked processus postorbitalis deforms the dorsal edge of the zygomatic process, at the junction between the jugal and the squamosal. Its tip, located on the latter bone, has a rugose aspect. Most of the temporal fossa elements are not preserved and it is therefore impossible to consider the shape and relations of the foramina sphenorbitale and rotundum. The area between the temporal and nuchal crests is depressed, forming a deep gutter. The external auditory pseudo-meatus is partly closed ventrally. The posterior side of the processus zygomaticus is flat in lateral view (no posterior groove). The occipital side is inclined up- and forward, with a very salient nuchal tubercle (although small, i.e., not extended on a wide area), determining a diamond-shape to the skull in dorsal view. The occipital condyles are oriented in the same axis as the skull in lateral view. The posterior tip of the tooth row reaches the posterior third of the skull. The pterygoids are not preserved, as most of the basicranium, vomer, and basal foramina. The skull is brachycephalic (interzygomatic width/total length ∼0.57; Table 1). As observable in dorsal view, the nasals have a sharp tip. They are long and unfused, fully separate by a deep groove from tip to tip. There were no lateral nasal horns, but a small median nasal horn, as unambiguously shown by the presence of axial vascularised rugosities in the anterior quarter of the nasal bones. In contrast, the frontal bones have a smooth aspect, thus indicating the absence of a frontal horn. The orbits were not projected laterally. The zygomatic arches are 1.51 times wider than the frontals. From this frontal ambitus, run posteriorly two straight and smooth frontal crests, getting closer by the parietals (minimum distance = 30 mm; Table 1), and then abruptly diverging and forming an occipital crest that is concave posteriorly. The transition from the maxilla to the processus zygomaticus maxillary is progressive, with no brutal inflection. The articular tubercle of the squamosal is smooth (in lateral view) and straight (in sagittal view). The right processus postglenoidalis forms a rounded right dihedron in ventral view. The foramen postglenoideum is remote from the latter. The left one is not preserved. The occipital side is wide and, accordingly, the processus posttympanicus and the processus paraoccipitalis are distant. The former is poorly developed, while the latter is very long, slender, and vertical. The foramen magnum is not preserved well enough to allow any observation. The occipital condyle has a median ridge but no medial truncation.

### Mandible

In lateral view, the symphysis is upraised, with an angular ventral profile determined by two successive inflections. The foramen mentale is widely open and located below p2 (left) and p3 (right). The corpus mandibulae is low, with a straight ventral border (Table 2). It is getting gradually higher to the mandibular angle, smooth, rounded and hugely developed. There is a shallow vascular incisure. The ramus is low, with a posterior border that is oblique up- and forward and a vertical anterior border. The processus coronoideus is high, tapering dorsally, and somewhat concave posteriorly. The condyloid process is high and sharp-edged, separate from the latter by a deep mandibular notch. In dorsal view, the symphysis is massive, well developed anteroposteriorly and narrow, with i2s and lateral edges parallel and two circular alveoli for small i1s. The posterior border of the symphysis is located between the trigonids of p3. The tooth rows are more parallel than the bodies (Fig. 4), which widely diverge posteriorly. The spatium retromolare is wide on both sides. The mylohyoid sulci are present but very shallow. The foramen mandibulare opens below the teeth-neck line. Dental material

**Fig. 4.**
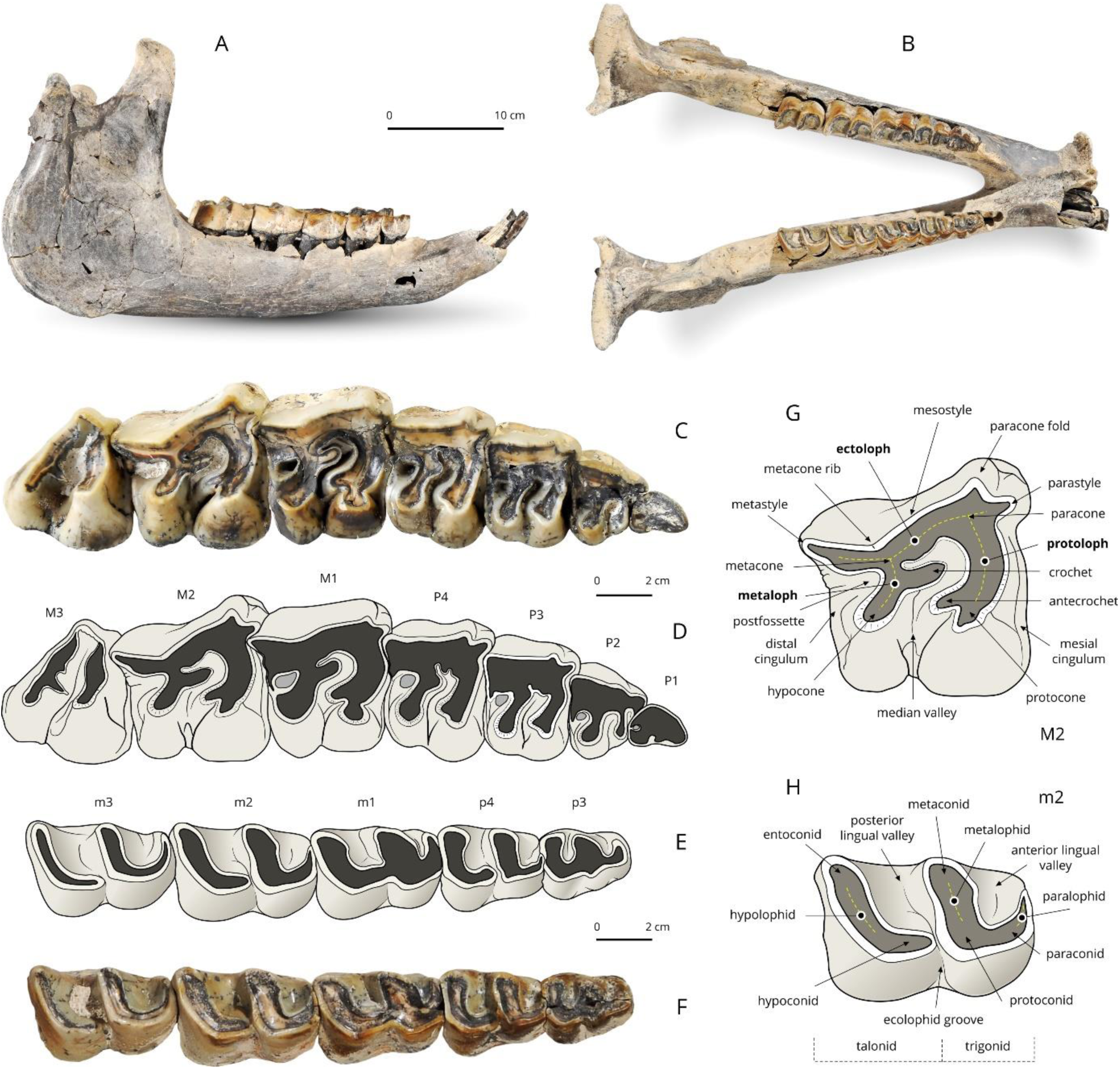
*Brachydiceratherium shanwangense* (Wang, 1965) from Tagay, Baikal Region, Russia, late early Miocene. Mandible and dental material. A, B – Mandible in right lateral (A) and occlusal views (B); C, D – Right upper cheek teeth (D1–M3) in occlusal view: photograph (C) and interpretative sketch (D); E, F – Right lower cheek teeth (p3–m3) in occlusal view: photograph (E) and interpretative sketch (F); G, H – Dental terminology used for rhinocerotid upper tooth (G) and lower tooth (H).

**Table 2.** Right mandibular measurements of *Brachydiceratherium shanwangense*, from Tagay, early Miocene of Eastern Siberia, in mm. 1, Maximal length; 2, Length without the symphysis; 3–8, Heights of the corpus mandibulae, between p2-p3, p3-p4, p4-m1, m1-m2, m2-m3, and behind m3, respectively; 9 and 10, Transverse diameters of the corpus mandibulae, between p4-m1 and behind m3, respectively; 11, Antero-posterior length of the symphysis; 13, Antero-posterior diameter of the ramus (at the level of the occlusal line); 14, Transverse diameter of the articular condyle; 15, Height of the articular condyle; 16, Height of the coronoid process. Numbers coincide with measurements as defined and illustrated by Guérin (1980, fig. 1, table 3).

| 1 | 2 | 3 | 4 | 5 | 6 | 7 | 8 | 9 | 10 | 11 | 13 | 14 | 15 | 16 | L. diast. |
| --- | --- | --- | --- | --- | --- | --- | --- | --- | --- | --- | --- | --- | --- | --- | --- |
| 467.4 | 136.8 | 68 | 68 | 70.7 | 75 | 76.4 | 77.7 | 38.1 | 41.3 | 118.2 | 136.8 | 96.2 | 203.3 | 229.2 | 51 |

The dental formula is 1-0-4-3/2-0-3-3. No decidual dentition is known.

### Upper dentition (Fig. 4, Table 3)

The first upper incisors are not preserved, but straight and sagittally-elongated alveoli point to an oval cross section for them (as usual in teleoceratines). There are no I2, I3, or C. The premolar series is short with respect to the molar series (LP3-4/LM1-3*100=48.7; Lp3-4/Lm1-3*100=45.8), which is further highlighted by the small size of P2 and p2. The enamel is thick, wrinkled and corrugated, and partly covered with a thin layer of cement. Teeth are low crowned, with roots partly joined. The labial cingulum is absent on the upper cheek teeth. A thick paracone fold is present on P2-M3, vanishing with wear on P2-M1 and marked until the neck on M2-3. There is no metacone fold or mesostyle on the upper cheek teeth. Short and wide crochet is present on P3-4 (always simple), but absent on P2. There is no metaloph constriction on P2-4. The lingual cingulum is absent on all upper cheek teeth, except for a small tubercle on the anterolingual base of the hypocone on both P4. The postfossette forms a small and deep isometric pit. The antecrochet is getting stronger backward, from absent on P2 and short on P3-4 to very elongate on M1-3. The first upper cheek tooth is most likely a persistent D1: it is much more worn than other teeth and the enamel is also much thinner. It is preserved on the right side and its presence is further attested by two alveoli on the left side (heart-shaped anterior root; peanut-shaped posterior root). It has a sharp anterolingual cingulum, a straight lingual edge, and rounded posterior and labial edges. P2-4 are fully molariform (bilophodont, with an open lingual valley). On P2, the metaloph is transverse labially, but curved posterolingually due to the position of the hypocone. The latter is much more developed than the protocone. The protoloph is thin but continuous and transversely oriented. There is no medifossette on P3-4, but a short crista on P4 and on P3 (mostly wiped out by wear on P3). The protocone is constricted anteriorly on P3-4. The metaloph forms a dihedron on P3-P4, with the crochet as a tip and the hypocone located posterior to the metacone. The protoloph is complete and continuous and there is no pseudometaloph on P3. The metacone is not constricted or isolated on P3-4. The crochet is long and sagittal on M1-M3, with a rounded tip on M1, and a sharp tip on M2-3. There is no crista, medifossette, or cristella on upper molars. The lingual cingulum is restricted to a small pair of tubercles on M2s and a smooth ridge on the hypocone of M3. The protocone is strongly constricted on M1-3, and trefoil shaped on M3. The parastyle is short and sagittal on M1-3; the paracone fold is very salient on M2 and especially on M3. The metastyle is very long on M1-2. The metaloph is almost as long as the protoloph on M1-2. In lingual view, the protocone is increasingly developed sagittally from M1 to M3. A deep groove carves the anterolingual side of the hypocone on M2, and a shallower one is observed on M1. The ectoloph is straight on M1 and concave on M2. The antecrochet and the hypocone are close but separate on M1-2. There is no lingual groove on the lingual side of M2. The posterior cingulum is complete on M1-2 and the postfossette is still narrower and deeper than on premolars. The right M3 has a triangular outline in occlusal view, with a straight ectometaloph (the left M3 is not preserved). The protoloph is transversely developed. There is no posterior groove on the ectometaloph and the labial cingulum is restricted to a low and smooth spur covering the lingual third of the former.

**Table 3.** Dental measurements of *Brachydiceratherium shanwangense*, from Tagay, early Miocene of Eastern Siberia, in mm. Abbreviations: H, crown height; L, length; W, width.

|  |  | P1 | P2 | P3 | P4 | M1 | M2 | M3 | p2 | p3 | p4 | m1 | m2 | m3 |
| --- | --- | --- | --- | --- | --- | --- | --- | --- | --- | --- | --- | --- | --- | --- |
| Left | L |  | 25.1 | 31.5 | 35.3 | 47 | 50.5 |  |  | 28 | 31.8 | 38.6 | 43.8 | 43.9 |
|  | W |  | 31.5 | 41.1 | 47.7 | 51.7 | 54.2 |  |  | 21.8 | 25.2 | 27.6 | 29.8 | 30.5 |
|  | H |  | 17.1 | 21.7 | 28.3 | 30.5 | 37.5 |  |  | 18.2 | 23.2 | 28.8 | 30.2 | 29.8 |
| Right | L | 19.3 | 27.2 | 29 | 35.5 | 47.1 | 50.6 | 42.1 |  | 29.8 | 32 | 37.7 | 44.1 | 45.4 |
|  | W | 14.5 | 31.9 | 40 | 47.6 | 51.4 | 56.2 | 46.1 |  | 21.9 | 27.2 | 28.1 | 30 | 30.3 |
|  | H | 8.8 | 18 | 21.1 | 27 | 29.7 | 37.6 | 37.8 |  | 19.8 | 23 | 24.8 | 28.7 | 27 |

### Lower dentition (Fig. 4, Table 3)

There are small circular alveoli for both i1s, between the i2s, in the symphyseal part of the dentary but the shape of the concerned teeth is unknown. The presence of a short p2 is attested by three closely-appressed alveoli on the right side (area unpreserved on the left side), but no d1 or p1 was present, as attested by the sharp ridge running anterior to p2’s alveoli. There are no vertical rugosities on the ectolophid of p3. On the lower cheek teeth, ectolophid grooves are developed (U-shaped) and vanishing before the neck, trigonids are rounded and forming a right angle in occlusal view, metaconids and entoconids are unconstricted. The bottom of the lingual valleys is V-shaped in lingual view on lower premolars. On lower premolars, the lingual cingulum is restricted to a low ridge continuing the anterior cingulum on the trigonid of p3s, and the labial cingulum consists of a small edge obtruding the ectolophid groove on p4s. Lower molars lack a lingual cingulum but a small cingular ridge partly obtrudes the ectolophid groove. The hypolophid is oblique in occlusal view on m1-3. There is no lingual groove on the entoconid of m2-3. The posterior cingulum of m3 forms a low, horizontal, and transversely-elongated ridge.

### Poscranial skeleton (Tables 4, 5)

#### Atlas

The atlas is wide and short sagittally. In dorsal view, the transverse processes (partly broken) and the alar notches are developed and the axis-facets are concave. In anterior view, the rachidian canal has a bulb-like outline. The occipital condylar facets are kidney-like. The foramen vertebralis cuts across the anterior third of the dorsal surface on both sides and it is continued by a shallow groove laterally (for the vertebral artery). In posterior view, the foramen transversarium is present, wide and partly hidden by the lateral expansion of each axis-facet (Fig. 5, A).

**Fig. 5.**
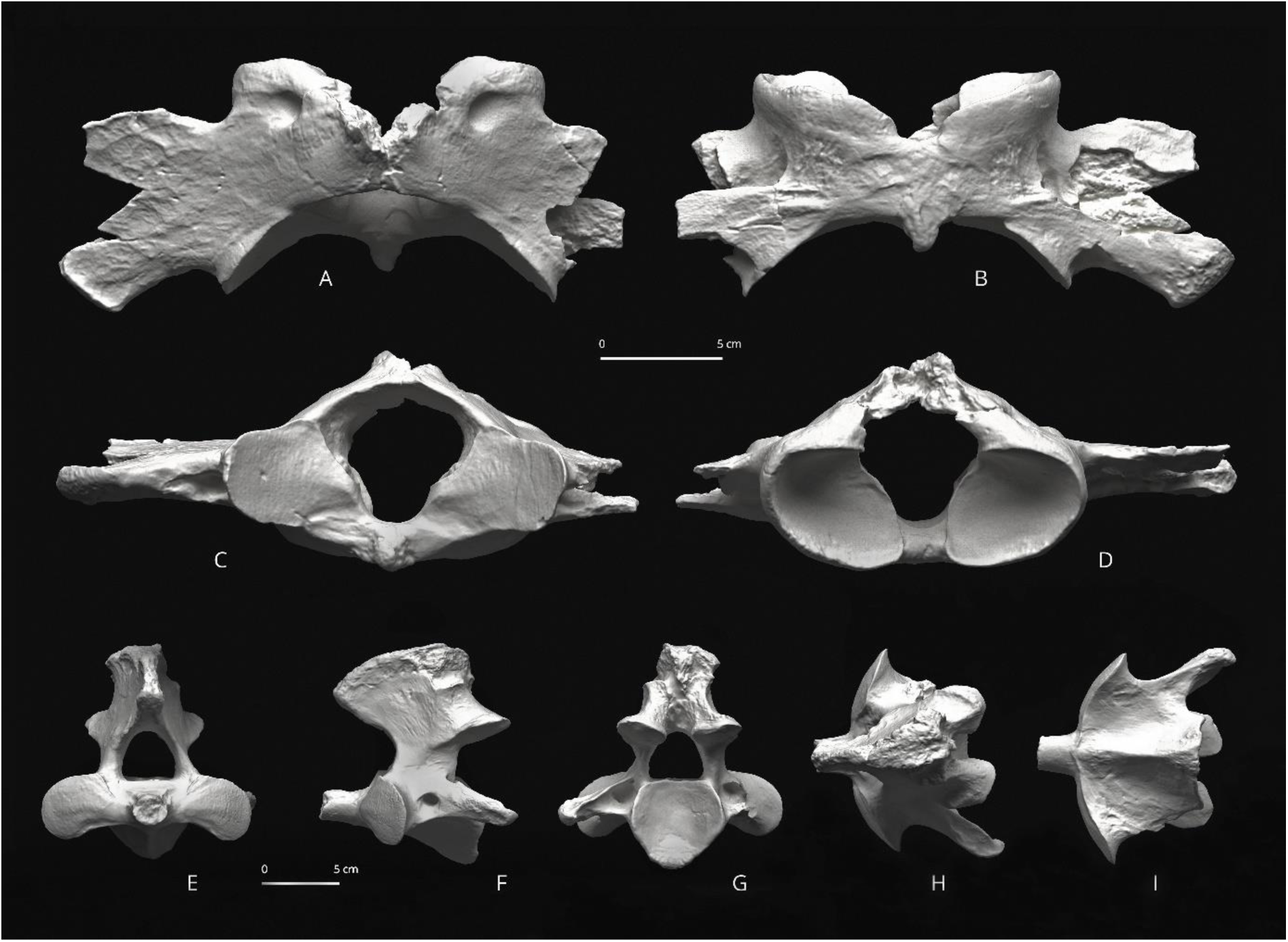
*Brachydiceratherium shanwangense* (Wang, 1965) from Tagay, Baikal Region, Russia, late early Miocene. A-D – Atlas in dorsal (A), ventral (B), cranial (C) and posterior views (D); E-I – Axis in anterior (E), left lateral (F), posterior (G), dorsal (H), and ventral views (I).

**Table 4.**
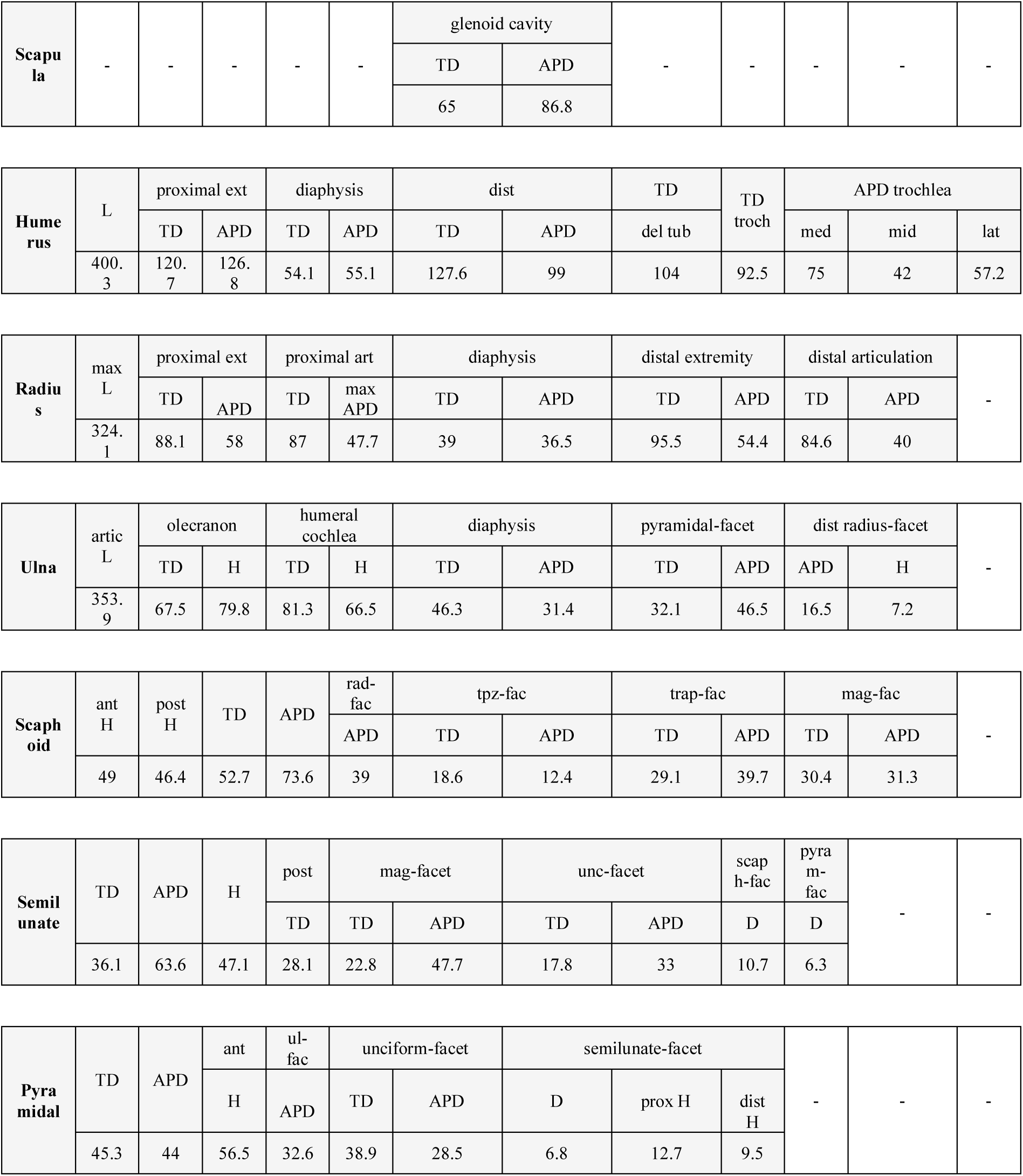
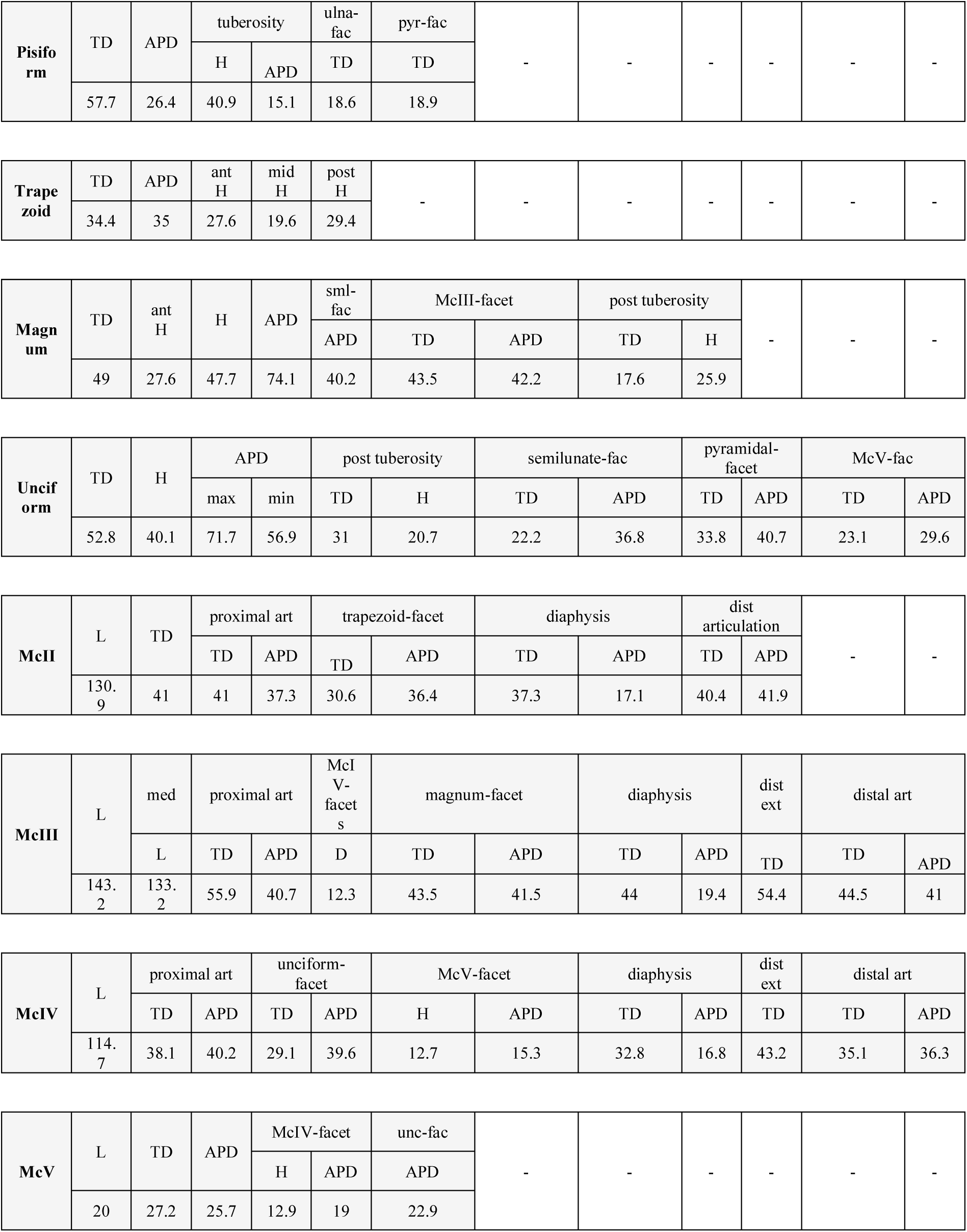
Postcranial measurements of *Brachydiceratherium shanwangense*, from Tagay, early Miocene of Eastern Siberia, in mm. Forelimb bones. Abbreviations: ant, anterior; APD, antero-posterior diameter; art, articulation; artic, articular; D, distance; del, deltoid; dist, distal; ext, extremity; H, height; L, length; lat, lateral; mag, magnum; max, maximal; med, medial; mid, middle; post, posterior; pyr, pyramidal; rad, radius; sml, semilunate; TD, transverse diameter; tpz, trapezium; tpzd, trapezoid; troch, trochlea; tub, tuberosity; unc, unciform.

**Table 5.**
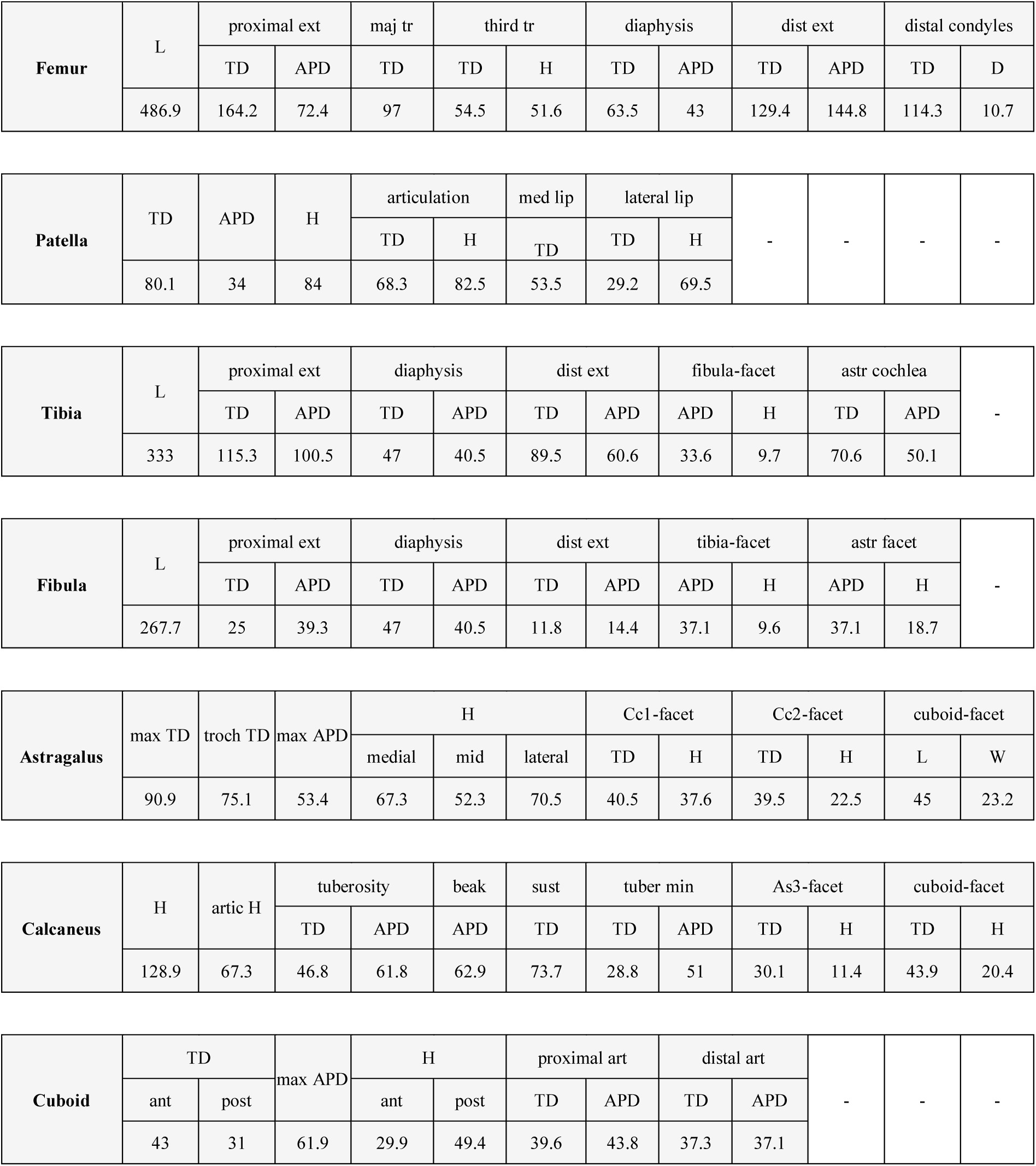
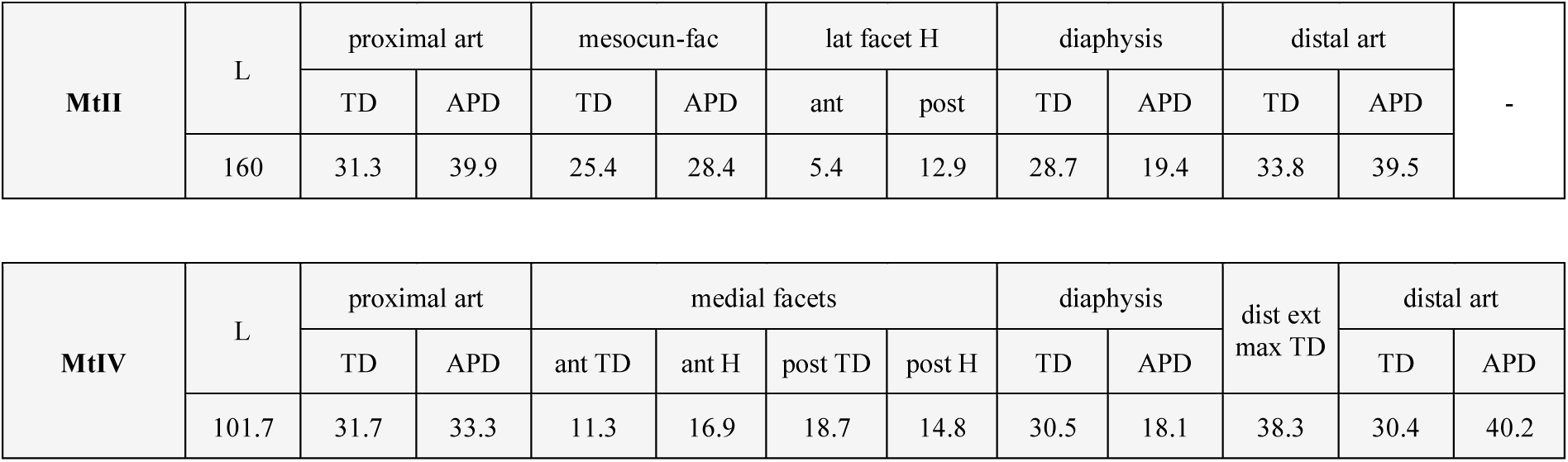
Postcranial measurements of *Brachydiceratherium shanwangense*, from Tagay, early Miocene of Eastern Siberia, in mm. Hind limb bones. Abbreviations: ant, anterior; APD, antero-posterior diameter; art, articulation; artic, articular; As3, astragalus-3 (sensu Heissig, 1972); astr, astragalus; Cc1, calcaneus-1 (sensu Heissig, 1972); Cc2, calcaneus-2 (sensu Heissig, 1972); D, distance; dist, distal; ext, extremity; H, height; L, length; lat, lateral; maj, major; max, maximal; med, medial; mesocun, mesocuneiform; mid, middle; min, minimum; post, posterior; sust, sustentaculum; TD, transverse diameter; tr, trochanter; troch, trochlea; tuber, tuberosity.

#### Axis

The axis is stocky, with thick and cylindric dens and tear-shaped atlas-facets (convex transversely) on the prezygapophyses. The spinous process is thick and carinated. The foramen vertebrale is large and subtriangular. The postzygapophyses have wide and circular facets for the first thoracic vertebra, forming a ∼45° angle with the horizontal line. The centrum is very long anteroposteriorly, with a pentagonal outline in posterior view (Fig. 5, B). Most thoracic vertebrae are preserved. They are massive, with heart-shaped centrums, and stocky transverse processes. The dorsal spines are slender and oblique (45° with the vertical line), with a length reaching up to 250% of the centrum height for the T4-6.

#### Scapula

The scapulae are partly preserved. They are elongated dorsoventrally, notably due to their anteroposterior narrowness (H/APD = ∼0.50). The scapular spine is straight, much developed and with an extremely salient tuberculum bent caudally. There is no pseudo-acromion. The tuberculum supraglenoidale is well distinct from the cavitas glenoidalis. The medial border of the cavitas glenoidalis is straight, determining a semi-circular outline in ventral view.

#### Humerus

Both humeri are almost complete (Fig. 6, A-E). The humerus is a slender bone, with a straight diaphysis. The trochiter is high, with a smooth and rounded outline. The caput humeri is wide and rounded, with a rotation axis forming a 40° angle with the vertical line. The deltoid crest is elongated, almost reaching the mid-bone. The deltoid tuberosity is not much salient. The fossa radii is wide and shallow. The fossa olecrani is higher than wide. The distal articulation is egg-cup shaped (sensu Antoine, 2002, 2003), without marked median constriction. The trochlea is half-conical and the capitulum humeri is half-cylindrical. There is no synovial fossa (“trochlear scar”) on the anterodorsal edge of the trochlea. The lateral epicondyle is elongated dorsoventrally and its ventral border ends dorsal to the capitulum humeri, lacking a distal gutter.

**Fig. 6.**
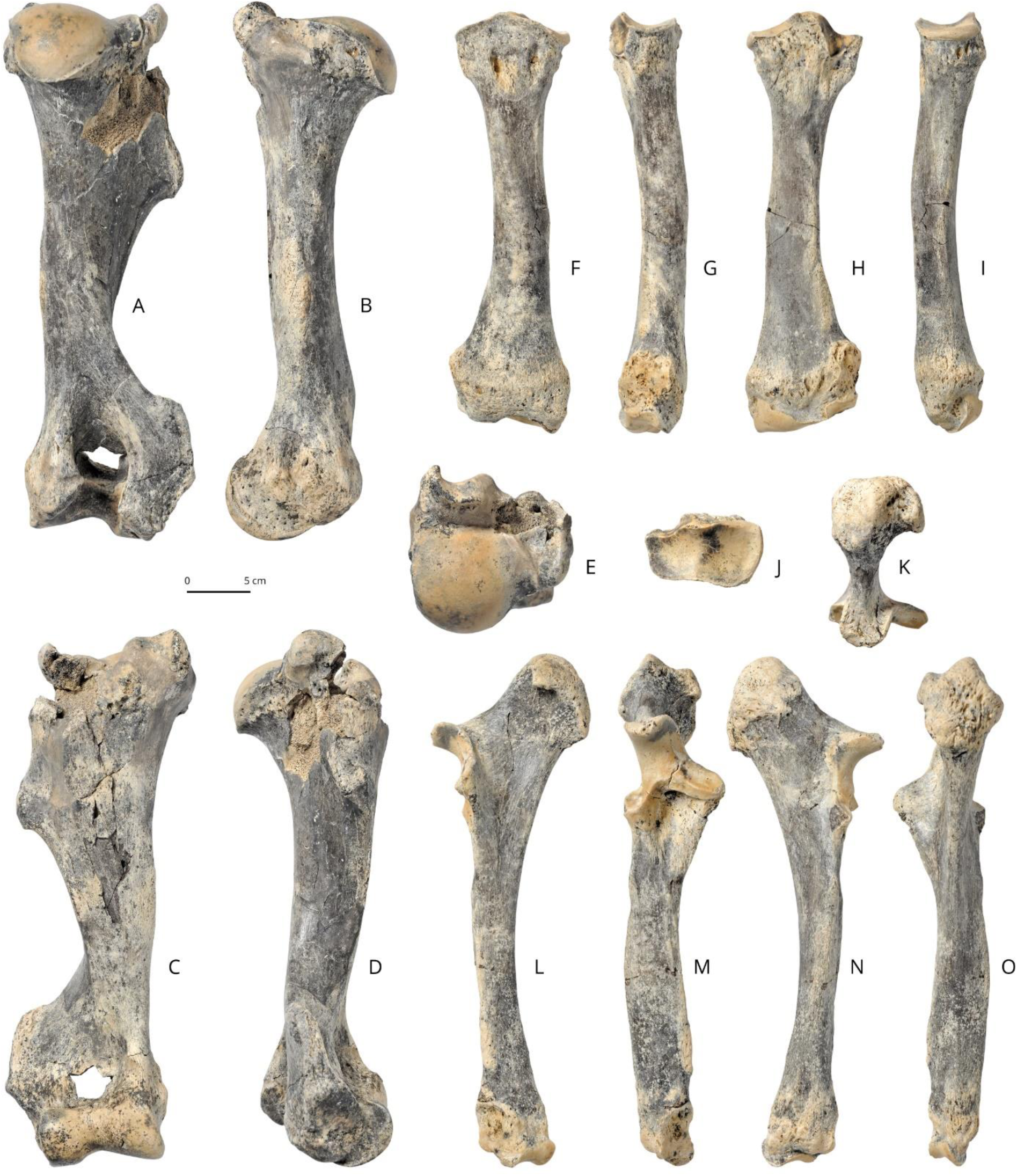
*Brachydiceratherium shanwangense* (Wang, 1965) from Tagay, Baikal Region, Russia, late early Miocene. Long bones of the right forelimb. A-E –humerus in posterior (A), medial (B), anterior (C), lateral (D), and proximal views (E); F-J – radius in anterior (F), lateral (G), posterior (H), medial (I), and proximal views (J); K-O – ulna in proximal (K), medial (L), anterior (M), lateral (N), and posterior views (O).

#### Radius

The two bones are complete and undistorted (Fig. 06, F-J). The anterior border of the proximal articulation is straight in dorsal view but convex in anterior view. The radius is slender, with a distal extremity larger than the proximal one in anterior view. The diaphysis is quite slender, especially in its proximal half. It has a straight medial border in anterior view, but it is posterolaterally concave, which determines a wide spatium interosseum brachii when the ulna is in anatomical connection. The proximal ulnar facets are fused on both sides. The insertion of the m. biceps brachii is wide but shallow, with two small pits. Ulna and radius are independent, apart from the proximal and distal articular areas. On the anterodistal part of the diaphysis, the gutter for the m. extensor carpi is not marked at all. There is only one distal facet for the ulna on the lateral side of the bone. The posterior expansion of the scaphoid-facet is high, forming a right-angled rectangle. There is a wide pyramidal-facet on the distal articulation.

#### Ulna

The bone is sturdy, with a long and heavy olecranon, the tip of which is wide and diamond shaped (Fig. 6, K-O). The diaphysis is straight, triangular in cross-section and as robust as the radius shaft. It forms a *∼*135° angle with the olecranon in lateral view. The humeral facet is saddle-shaped. The proximal radio-ulna facets form a continuous pad, with a wide medial strip and a high triangular lateral facet. A smooth but salient anterior tubercle is located above the distal end of the bone. There is neither a second distal radius-facet on the medial side of the diaphysis nor semilunate-facet on the distal side. The almond-shaped distal radius-facet is separate proximally from a salient horizontal ridge by a deep and rugose depression. The pyramidal-facet is concavo-convex, with a quarter-circle outline in distal view.

#### CARPUS

The carpus is very low and massive, especially with respect to slender stylopodial and zeugopodial elements (Fig. 7). All carpals have salient tubercles on the anterior aspect of the bones. The right hand is more complete than the left one.

**Fig. 7.**
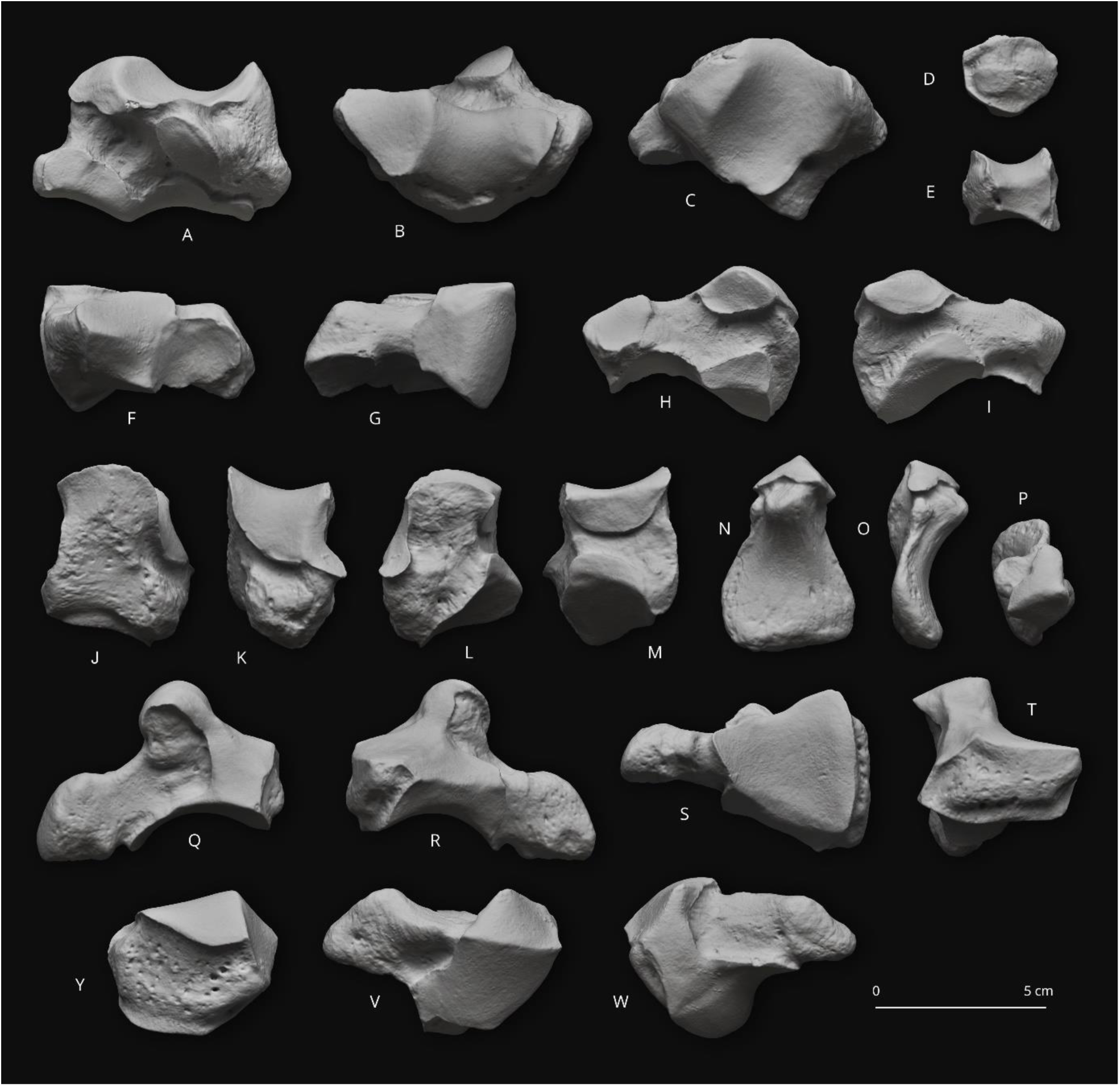
*Brachydiceratherium shanwangense* (Wang, 1965) from Tagay, Baikal Region, Russia, late early Miocene. Carpal bones. A-C – left scaphoid in posterior (A), proximal (B), and distal views (C); D-E – right trapezoid in anterior (D), and distal views (E); F-I – left semilunate in distal (F), proximal (G), medial (H), and lateral views (I); J-M – left pyramidal in anterior (J), lateral (K), posterior (L), and medial views (M); N-P – right pisiform in anterior (N), lateral (O), and proximal views (P); Q-T – right magnum in lateral (Q), medial (R), distal (S), and anterior views (T); Y-W – right unciform in anterior (Y), proximal (V), and distal (W).

#### Scaphoid

The scaphoid is low and massive, with equal anterior and posterior heights (Fig. 7, A-C). The proximal radial facet is diamond shaped in proximal view. The posteroproximal semilunate facet is strongly distinct. It is oval, wide, and separated from all other facets. A deep depression hollows the lateral side between the semilunate-facets. The anterodistal semilunate-facet is nearly flat and crescent shaped. The magnum-facet is concave in lateral view. The trapezium-facet is smaller than other distal facets, but it forms a wide triangle, separated from the trapezoid-facet by a smooth ridge.

#### Semilunate

The bone is compact. In proximal view, the anterior facet only contacts the radius, whereas the wide posteromedial facet is for the scaphoid (Fig. 7, F-I). The anterior side is smooth (not keeled or carinated), with a sharp distal tip. On the lateral side, both pyramidal-facets are closely appressed. The proximal one is almond shaped and the distal one is comma like. The posterior tuberosity is short. Most of the distal side is articulated, medially with the magnum and laterally with the unciform.

#### Pyramidal

The bone is almost cubic. The proximal side is square shaped, with a saddle-shaped ulna-facet (Fig. 7, J-M). The semilunate-facets are sagittally elongated, with a half-oval outline for the proximal one and a crescent-like shape for the distal one. The pisiform-facet is comma shaped, with a concave sagittal profile and it overhangs a strong lateral tuberosity. The distal facet, for the unciform, forms a right isosceles triangle with rounded angles. There is no magnum-facet.

#### Pisiform

The right pisiform is short, high, and spatulate, with large and triangular radius- and pyramidal-facets (Fig. 7, N-P). Both facets are separated by a sharp ridge and form a right angle. There is no strong constriction separating the thick body and the articulated part. The medial edge of the body is straight and vertical.

#### Trapezium

The right trapezium is preserved. It is a small proximo-distally flattened bone with a circular outline in proximal view. The proximal side is almost entirely occupied by a wide pentagonal scaphoid-facet (compatible with the large-sized trapezium-facet on the scaphoids). The latero-distal side bears a trapezoid-facet with a right-triangled shape, overhanging a deep pit. All other sides have a rugose aspect and they are devoid of articular facets.

#### Trapezoid

Only the right trapezoid is documented (Fig. 7, D-E). It is wider than high, almost cubic. Only the anterior and posterior sides (oval and pentagonal in shape, respectively) are free of articular surfaces. The proximal side, saddle shaped and tapering posteriorly, responds to the scaphoid. In medial view, the trapezium-facet is restricted to the posterior half, with a deep insertion pit located close to the anterior edge. The lateral facet is al low rectangle for the magnum. The distal side is weakly concavo-convex, and it consists of a pentagonal McII-facet.

#### Magnum

The magnum has a very low anterior aspect, with a subrectangular outline and a salient horizontally-elongated median pad (Fig. 7, Q-T). The proximal border is straight in anterior view. In medial view, the anteromedial facets are in contact throughout their whole length (no anterior groove). In lateral view, the dorsal pulley for the semilunate forms a low-diameter half circle, further determining a question mark proximal profile. The distal facet is wide and tapering posteriorly. The posterior tuberosity is broken on the left magnum, and it is very short on the right specimen.

#### Unciform

The bone is compact, with a posterior tuberosity wide and much developed sagittally (Fig. 7, Y-W). The anterior side is wide and low, with a pentagonal outline and a maximum height on its lateral tip. The proximal side has two anterior facets flat transversally and convex sagittally, separated by a sharp sagittal edge. The medial one, triangular, is for the semilunate while the lateral one, diamond shaped, is for the pyramidal. The latter has a wide posterolateral expansion joining the lateral edge and the McV-facet (located on the distal side) on the right unciform. This part is broken on the left one. From the medial tip, the distal and distolateral sides have three contiguous facets, responding to the McIII (small and quadrangular), McIV (bulb-shaped), and McV (oval and deeply concave sagittally), respectively. They are only separated by smooth sagittal grooves. The McV-facet is oblique, which could suggest the presence of a functional McV (see Antoine, 2002, 2003; Boada-Saña et al., 2008).

#### METACARPUS

The hand and pes have a mesaxonian Bauplan. Although no McV is preserved, the hand was probably tetradactyl, as hypothesised by the vertical facet on the McIV (see above). The metapodials have salient insertions for the m. extensor carpalis. Their shafts are robust (wide transversally and flattened sagittally), with neither distal widening nor clear shortening (no brachypody; see Antoine, 2002). The insertions for the m. interossei are long, reaching the mid-shaft on all available metapodials. The intermediate reliefs do not reach the anterior aspect of the distal articulation on metapodials. The intermediate relief is moderately high and quite sharp on the McIII, but low and smooth on medial and lateral metapodials.

#### McII

In proximal view, the proximal side consists of a large tear-shaped trapezoid-facet medial to a narrow sagittally-elongated and strip-like magnum-facet. In medial view, the trapezium-facet is large and comma shaped, higher in its posterior tip. In lateral view, the magnum-facet is a straight and low strip, separated from the McIII-facets over their length. The McIII-facets are fused into a curved strip with a shallow disto-median constriction. The distal articulation, for the phalanx 1, has a sub-square outline in distal view, with rounded anterior angles. Above it, is a wide and salient medial tuberosity (Fig. 8, A-E).

**Fig. 8.**
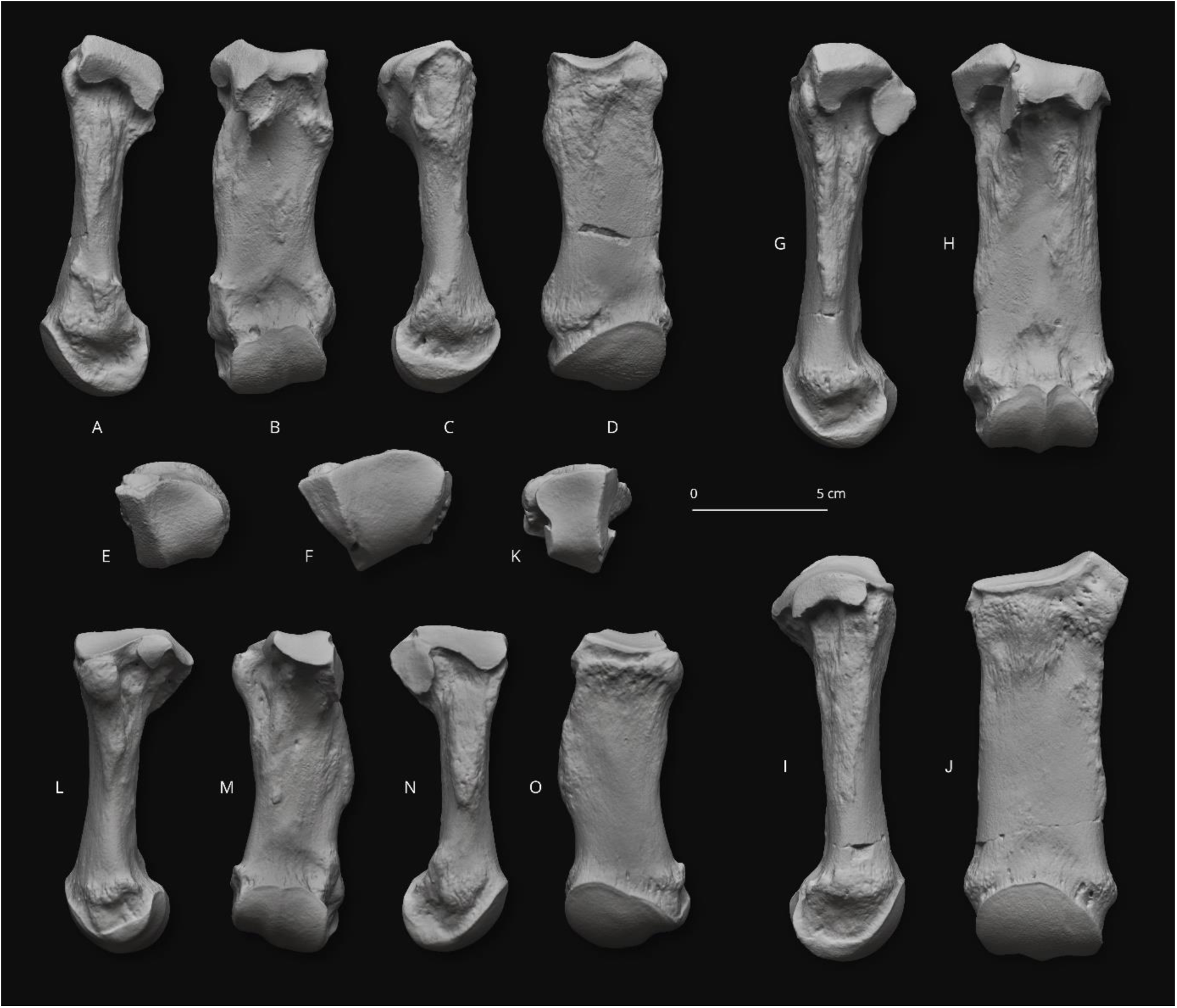
*Brachydiceratherium shanwangense* (Wang, 1965) from Tagay, Baikal Region, Russia, late early Miocene. Left metacarpal bones. A-E – second metacarpal in lateral (A), posterior (B), medial (C), anterior (D), and proximal views (E); F-J – third metacarpal in proximal (F), lateral (G), posterior (H), medial (I), and anterior views (J); K-O – fourth metacarpal in proximal (K), lateral (L), posterior (M), medial (N), and anterior views (O).

#### McIII

The bone has a straight shaft. The proximal side is dominated by a wide and pentagonal magnum-facet, contiguous to two narrow sagittal strip-shaped facets (medially for the McII and laterally for the McIV). In anterior view, the proximal side consists of a subvertical medial edge (McII-facet), a very wide magnum-facet, weakly-concave medially, and a much narrower, oblique and straight McIV-facet. The magnum-facet is almost invisible in anterior view. Indeed, its dorsal outline is not much convex in medial view. The McII-facets are broadly connected, forming a thick strip with a shallow constriction in its disto-median part. In lateral view, the anterior McIV-facet is low, elongated sagittally, and tear-shaped. It is disconnected from the oval posterior McIV-facet by a narrow but deep oblique groove. This articulated surface overhangs a deep circular depression. There is no postero-distal tubercle on the diaphysis. In distal view, the distal articulation is wide and subrectangular, with straight medial and lateral edges, rounded antero-medial and -lateral angles, and a m-like posterior edge, due to a low but sharp intermediate relief (Fig. 8, F-J).

#### McIV

The McIV is the shortest and most robust metapodial preserved. The shaft is concave laterally in anterior view. The proximal aspect is trapezoid, deeper than wide, with a narrow medial strip (the sagittally-elongated ‘anterior’ McIII-facet) and a wide unciform-facet. In proximal view, there is no postero-lateral pad, but a small anterolateral tubercle in front of the McV-facet. In medial view, the proximal McIII-facets are connected (right specimen) and form a right dihedron (L-shape), with a high posterior facet. The McV-facet is vertical, suggesting a functional McV, in good agreement with the orientation of the McV-facet on the unciform. In distal view, the distal articulation forms a quarter circle, with a posteromedial right angle. There is almost no intermediate relief on the McIV (Fig. 8, K-O).

#### Phalanges

Only three phalanges are preserved for the manus (left/right first phalanges and left second phalanx for the McII). They have strong interphalangeal insertions and tubercles. The phalanx 1 is low and massive, with a kidney-like proximal side (McII-facet, lacking a groove responding to the intermediate relief). The distal facet is oval and transversely transversally elongated. The phalanx 2 is still lower, with a proximal facet perfectly matching in shape the distal facet on the phalanx 1. The distal facet is slightly concave transversally and convex sagittally. Both facets have similar width and depth.

#### Coxal

The pubic bones and ischia are lacking on both sides but the ilia are well preserved. Dorsally, the iliac crest is regularly convex. The wing of the ilium is spatulated. The sacral tuberosity has a rounded triangular shape, with a rugose aspect. The coxal tuberosity, partly broken, was probably thick and high, also with a rugose aspect. The caudal gluteal line is smooth, with a concave outline (forming a semi-circular curve). The acetabulum has a subcircular outline.

#### Femur

The bone is quite slender, with a shaft straight in anterior view, concave anteriorly in lateral view, and compressed sagittally (Fig. 9, A-F). The anterior part of the trochanter major is high, but the caudal part is very low, i.e., much lower than the wide and hemispheric head. The fovea capitis is deep, low, and wide, with a triangular outline. The trochanter minor is elongated dorsoventrally. Its distal end reaches the mid-height of the third trochanter. The latter is developed, wider distally and with smooth lateral borders. The anteroproximal border of the patellar condyle is curved, with a medial lip much more developed and salient than its lateral counterpart. In lateral view, the medial lip of the trochlea and the diaphysis determine a broken angle (130°). In distal view, the anterior border of the patellar trochlea is convex medially and straight and transverse laterally. The tibial condyles are separate from the patellar trochlea by a narrow groove. The intercondylar fossa is deep and narrow. The medial condyle, with a diamond-shaped outline, is much more developed than the lateral one. The medial epicondyle is also more salient than the lateral epicondyle.

**Fig. 9.**
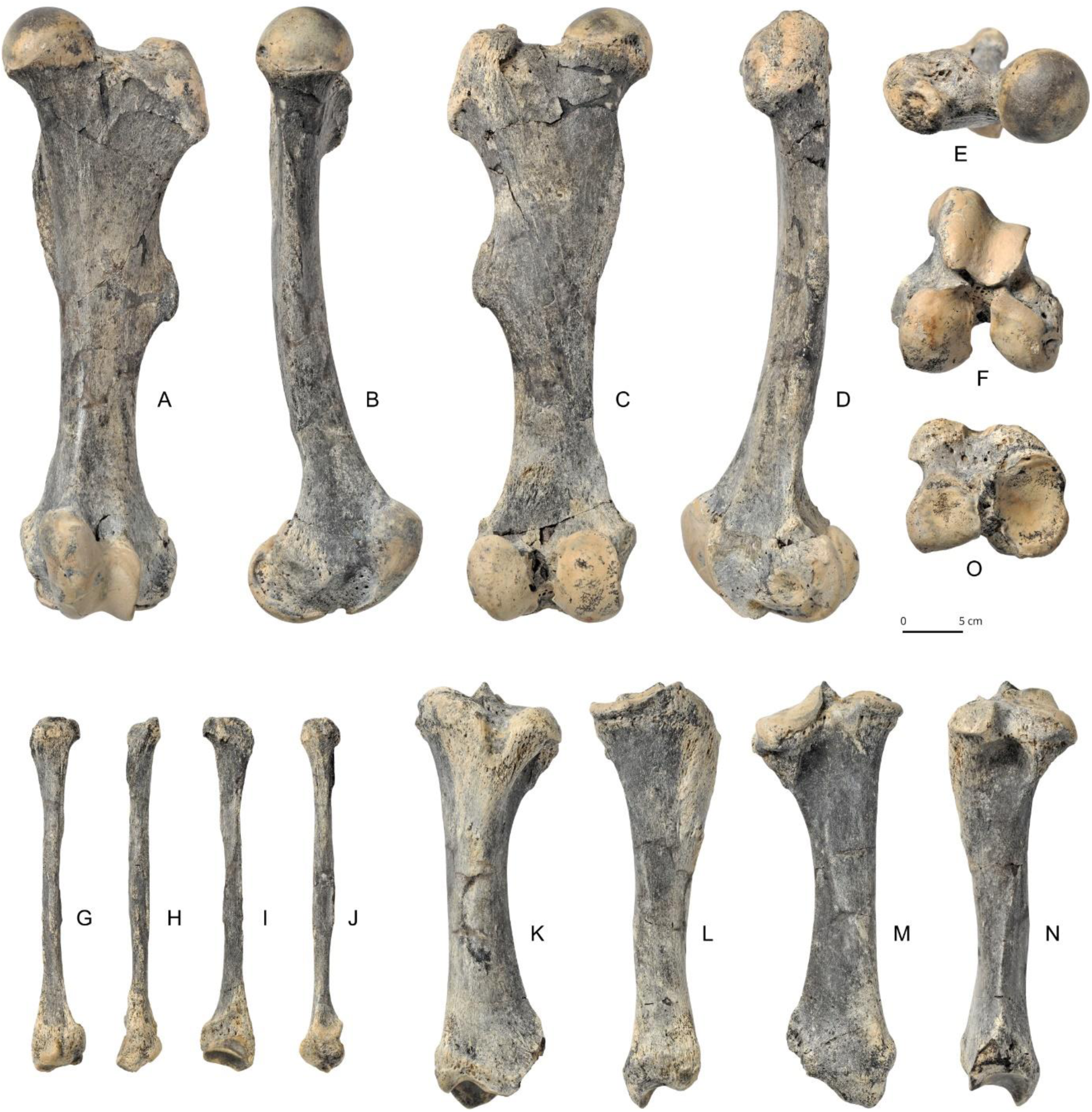
*Brachydiceratherium shanwangense* (Wang, 1965) from Tagay, Baikal Region, Russia, late early Miocene. Long bones of the left hind limb. A-F – femur in anterior (A), medial (B), posterior (C), lateral (D), proximal (E), and distal views (F); G-J – fibula in lateral (G), posterior (H), medial (I), and anterior views (J); K-O – tibia in anterior (K), medial (L), posterior (M), lateral (N), and proximal views (O).

#### Patella

The patella is massive, wider than high, and with a triangular and rugose anterior aspect. The medial border is straight and vertical. The posterior side, almost fully articulated, contacts the femoral cochlea, with a wide medial lip, triangular (wider distally), and a narrower trapezoid lateral lip. In vertical view, the latter lip is almost straight while the former is more concave. The most striking feature is the weak anteroposterior development of the bone with respect to other dimensions.

#### Tibia

The tibia is high and relatively slender, with heavy extremities (Fig. 9, K-O). The medial border of the diaphysis is strikingly straight in anterior view, which widely contrasts with the concave lateral border of the shaft. This impression is highlighted by the median half of the proximal articulation being much higher than the lateral one. The patellar ridge is thick and bulbous, with a rough surface. The patellar groove is deep, short dorso-ventrally, and regularly concave. The proximal peroneal articulation is located low on the tibia (no contact with the lateral femoral condyle). There is neither an anterodistal groove nor medio-distal gutter (for the tendon m. tibialis posterior). Tibiae and fibulae are independent, apart from articulated areas, thus determining a wide spatium interosseum cruris. The distal fibula-facet is low, elongated, and crescent shaped, overhung by a rugose triangular area. The posterior apophysis is low and rounded. In distal view, the outline is a trapezoid, wider than deep. The astragalar cochlea has two lips, the medial one being narrower and deeper and the lateral one wider and shallower.

#### Fibula

The diaphysis is straight and particularly slender, in sharp contrast with two thick ends (and the robustness of the tibia) (Fig. 9, G-J). The proximal end is nevertheless flattened sagittally, with a smooth proximal tibia-facet. The distal end is robust, with a deep laterodistal gutter for the tendon m. peronaeus, located posteriorly, immediately posterior to a huge tubercle. The distal fibula-facet is low, elongated sagittally, and crescent shaped. It is contiguous to a flat and rectangular astragalus-facet, oriented at ∼25° with respect to the vertical line.

PES. The pes is not completely known (Fig. 10). The naviculars, cuneiforms, MtIIIs, and most phalanges are not preserved. The metatarsals are shorter than the metacarpals.

**Fig. 10.**
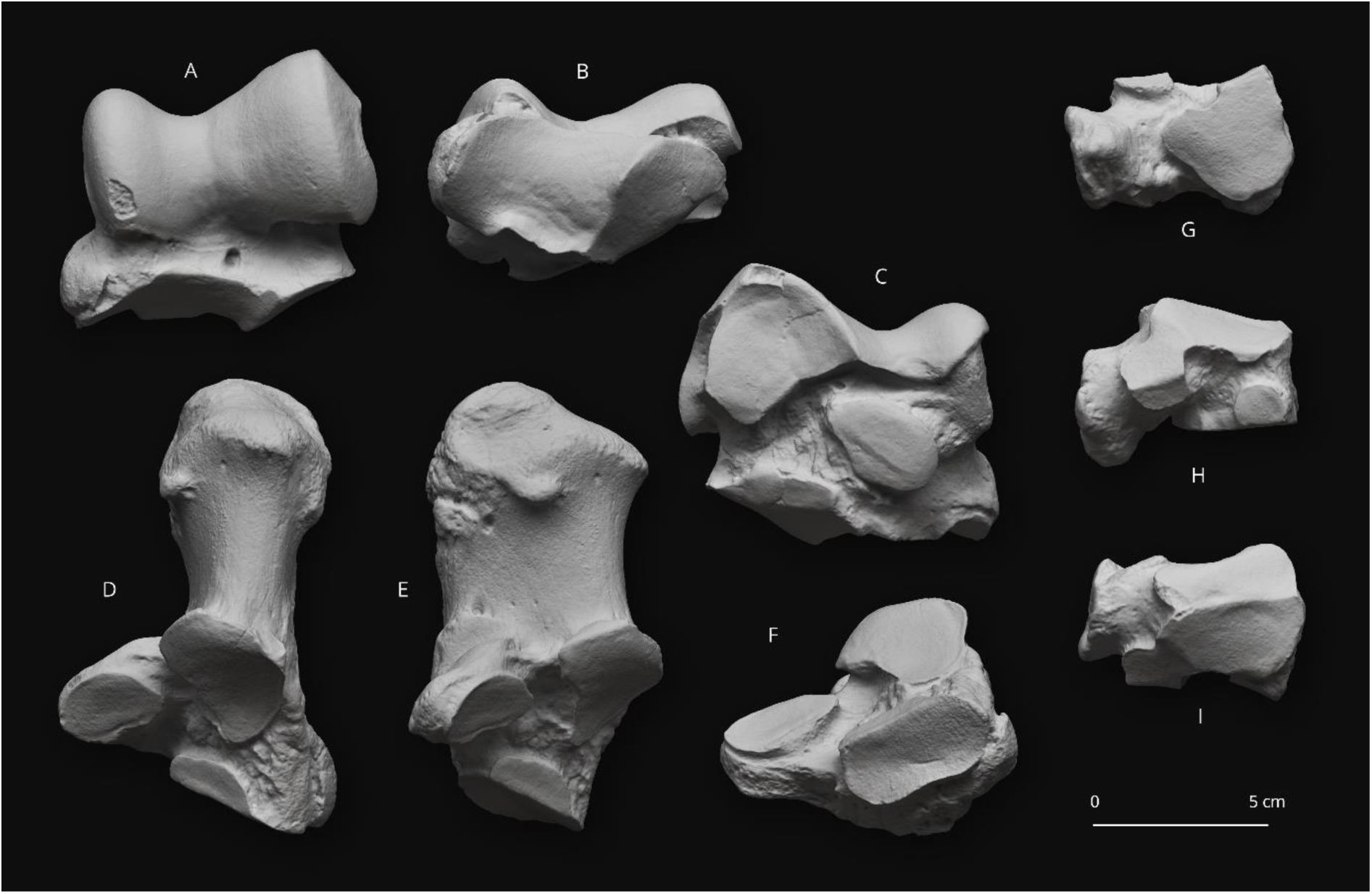
*Brachydiceratherium shanwangense* (Wang, 1965) from Tagay, Baikal Region, Russia, late early Miocene. Tarsal bones. A-C – left astragalus in anterior (A), distal (B), and posterior views (C); D-F – left calcaneus in proximal (D), medial (E), and anterion views (F); G-I – left cuboid in distal (G), lateral (H), and proximal views (I).

#### Astragalus

The astragalus is thick (APD/H = 0.76), wide and low (TD/H = 1.29). The fibula-facet is subvertical, wide and flat dorsoventrally (Fig. 10, A-C). The medial trochlear ridge is rounded, whereas the lateral one is sharper. The collum tali is very high (up to ¼ of the height), especially with respect to the general proportion of the bone. The caudal border of the trochlea is sinuous in dorsal view (with a falciform shape). There is no anterodistal trochlear notch, but a wide foramen for an insertion located distally to the concerned area, in the mid-collum tali. In anterior view, the distal border is deeply concave medially (navicular-facet) and straight and oblique laterally (cuboid-facet). The medial tubercle is low and rounded, but much projected medially. The distal articulation is not twisted with respect to the axis of the trochlea (<15°), in distal view. The calcaneus-facet 1 has a wide and very low, triangular laterodistal expansion. This facet is nearly flat in lateral view. The calcaneus-facets 2 (low oval) and 3 (tear shaped and low) are distinct and separate by a wide groove. In distal view, the distal articulation is much wider than deep, with a cuboid-facet particularly wide transversely. The posterior stop on that cuboid-facet is abrupt and prolongated medially by a similar transversely-elongated inflection on the navicular-facet.

#### Calcaneus

The calcaneus is robust, with a tuber calcanei massive and oval in posteroproximal view (Fig. 10, D-F). This tuber calcanei is strongly vascularised and rugged with salient muscle/tendon insertion areas, The tibia-facet is low, wide, and almond shaped, while the fibula-facet is round and oblique with respect to the vertical and sagittal lines. The astragalus-facet 1 is lozenge shaped in anterior view and almost flat. The facet 2 is oval, wider than high and flat. It is separate from the smaller and semi-oval facet 3. The sustentaculum tali is low and very wide. In lateral view, the cuboid-facet and the posterior border of the tuber form a right angle and the processus, at the level of the sustentaculum tali, is deeper (in terms of APD) than the tuber calcanei. The insertion for the m. fibularis longus forms a salient and rugose pad, but without sharp ridges. On the distal side, the cuboid-facet forms a transversely-elongated hexagon. It is mostly flat but concave in its mediodistal quarter.

#### Cuboid

The cuboid is compact, wide, and low (Fig. 10, G-I). In proximal view, the large articular surface is oval, slightly tapering posteriorly, and split into two equally-developed and sagittally-elongated facets. The astragalus-facet (medial) is separated from the calcaneus-facet (lateral) by a narrow and shallow groove. The anterior side is low and pentagonal in anterior view, with a sharp proximal tip. In medial view, there are four facets. The anteroproximal one is very low and crescent like (navicular-facet). Distally to it is a much larger semi-circular ectocuneiform-facet. The posteroproximal navicular-facet, broadly joining the proximal facet for the astragalus, has an 8-shaped outline. Contiguous to it, but distally, is a semi-circular posterodistal ectocuneiform-facet. The posterior tuberosity is short sagittally and narrow, but quite elevated: its acuminated distal tip is positioned much more distally than the distal articulation (MtIV-facet). The latter facet is flat and trapezoid, with larger anterior, medial, and posterior sides and a shorter lateral border. There is no MtIII-facet.

#### MtII

The bone is short and robust (Fig. 11, A-E). The proximal side, with a triangular outline (widening posteriorly), responds to the entocuneiform (posteromedial facet, pentagonal, and oblique), the mesocuneiform (proximal-most facet, wide and trapezoid), and to the ectocuneiform (wide strip-like facet oblique and tapering anteriorly). In lateral view, the MtIII-facets are vertical, with a large triangular anterior facet and a much lower, oval posterior facet. Both are widely connected. The shaft is straight and subcircular in cross section. The distal end is stocky and square in distal view. The distal articulation has almost no intermediate relief, even in its posterior aspect.

**Fig. 11.**
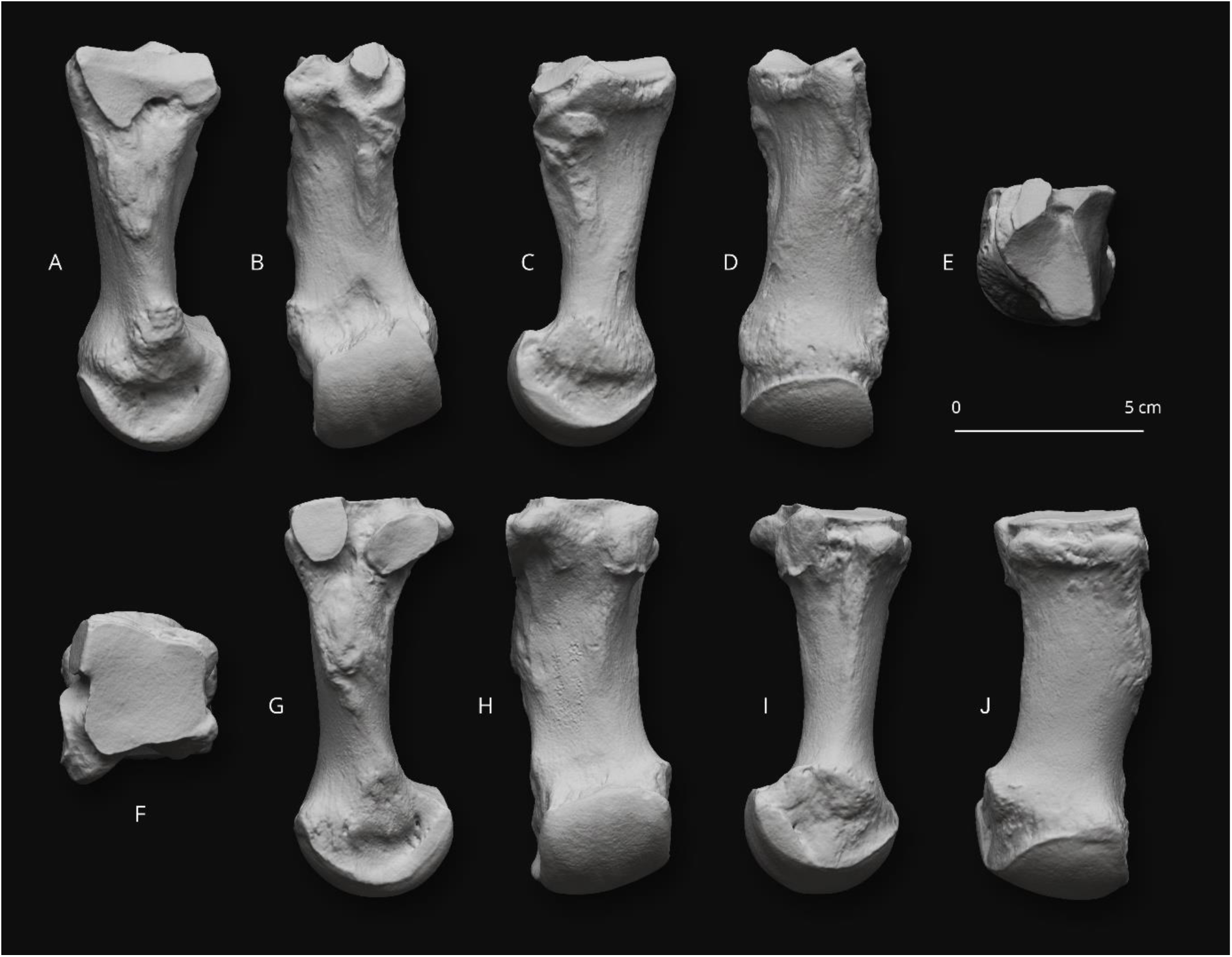
*Brachydiceratherium shanwangense* (Wang, 1965) from Tagay, Baikal Region, Russia, late early Miocene. Metatarsal bones. A-E – left second metatarsal in lateral (A), posterior (B), medial (C), anterior (D), and proximal views (E); F-J – right fourth metatarsal in proximal (F), medial (G), posterior (H), lateral (I), and anterior views (J).

#### MtIV

The bone is short and massive, with a heavy proximal end (Fig. 11, F-J). The proximal side is entirely occupied by a flat and sub-square cuboid-facet. There are two distinct proximal tubercles at the anterolateral and posterolateral angles, but no continuous pad. In medial view, there are two equally-wide MtIII-facets. The anterior one is located more dorsally, elevated and with a half-oval outline, connecting the proximal side. The posterior one is oval, isolated, and anteroventrally-posterodorsally elongated. The shaft is slightly concave laterally but straight in lateral view, with a strong laterodistal tubercle. The distal side is entirely articulated, deeper than wide (APD>TD), and lacking an intermediate relief. Only the lateral lip is slightly concave transversely in its posterior aspect.

#### Phalanges

Only the first phalanges for the MtII and MtIV are known. They have strong interphalangeal insertions and tubercles. There is no groove responding to the intermediate relief. The MtII phalanx 1 is almost cubic, with a circular and slightly biconcave proximal side (MtII-facet). The distal facet (phalanx 2) is kidney shaped. The MtIV phalanx 1 is as wide as but lower than the former phalanx. The proximal facet is kidney shaped and almost flat. The distal facet is oval and elongated transversely. In both phalanges, the distal facet is smaller than the proximal facet, but also slightly convex sagittally and concave transversely.

## COMPARISON

In the next pararaphs, the comparison will be organised following anatomical regions (skull, mandible, teeth, and postcranial skeleton) and we will use the generic assignments as supported by the phylogenetic analysis. The rhino from Tagay cannot be assigned to *Diaceratherium* (*sensu stricto*, see below) as it does not have teeth with wrinkled enamel, M1-2 with a short metaloph, M2 with a mesostyle, semilunates with a distal border of the anterior side rounded, or unciforms with a posterior expansion of the pyramidal-facet always present (all these features are diagnosing its type and only species, *D. tomerdingense*). It cannot belong either to *Brachypotherium*, the representatives of which have an occipital condyle without a median ridge, a mandibular symphysis very massive, a labial cingulum usually present on upper premolars and always present on upper molars, lower cheek teeth with a flat ectolophid, lower premolars with a lingual opening of the posterior valley U-shaped, p2 with a paraconid reduced, radius-ulnae with a second distal articulation, pyramidals with a distal semilunate-facet asymmetric, posterior facet always absent on the McII-McIII, and a fibula-facet oblique on the astragali. Contrary to representatives of *Prosantorhinus*, it has no sagittal fronto-parietal crest, posterior groove on the processus zygomaticus of the squamosal, or metacone fold on M1-2, but a constricted metaloph on M1 (diagnostic features of the genus). At last, it is differing from *Teleoceras fossiger* in a wide array of cranio-mandibular, dental, and postcranial features (e.g., foramen infraorbitalis above premolars, processus postorbitalis present on the zygomatic arch, occipital side inclined up-and frontward, low-crowned cheek teeth, crista always present on P3, atlas with a bulb-like rachidian canal cavity, scapula elongated, or navicular with a lozengic outline in vertical view).

On the other hand, in its general shape, proportions and anatomical features, the skull from Tagay closely matches that of *Brachydiceratherium shanwangense* (Wang, 1965) as recently described by Lu et al. (2021). It also resembles that of late Oligocene–early Miocene teleoceratines from Western Europe classically assigned to *Diaceratherium*, except for the type species (*D. tomerdingense*, for which only an isolated nasal bone is preserved). Within *Brachydiceratherium*, the arched dorsal profile, the short and slender premaxillae, the zygomatic arch (straight, oblique, with a marked posterodorsal angle, and an anterior tip starting progressively), and the processus paraoccipitalis long and narrow make it have the closest affinities with *Bd. shanwangense*, *Bd. aginense* (earliest Miocene; Répelin, 1917), and *Bd. asphaltense* (Becker et al., 2018). The only differences with the former do concern the tip of the nasal bones, pointing upward, having a small median bump (suggesting the presence of a terminal nasal horn) and a distolateral apophysis, and the stockier zygomatic arch as observed in the Tagay specimen. More specifically, the Tagay skull differs from that of *Bd. aginense* in having a processus postorbitalis on the frontal bone and a median ridge on the occipital condyle, but no posterior groove on the processus zygomaticus of the squamosal (Répelin, 1917). It is distinct from *Bd. lemanense* in possessing a low zygomatic arch, with a processus postorbitalis, a small processus posttympanicus and a well-developed processus paraoccipitalis (Lavocat, 1951), and from *Bd. asphaltense* in having closer fronto-parietal crests and a brachycephalic shape (Becker et al., 2018; Jame et al., 2019). In contrast, the shape of the processus zygomaticus, but also the presence of a small median nasal horn and of a concave occipital crest make is somewhat resemble *Bd. asphaltense*.

As for mandibular features, the Tagay jaw is also particularly resembling those of *Bd. shanwangense* and *Bd. aginense* among representatives of *Diaceratherium*, with an upraised symphysis (distinct from that of *Bd. lamilloquense* and of *Bd. aurelianense*), low corpus, vertical ramus and a deep and laterally-salient mandibular angle, with a shallow vascular incisure.

With respect to all other representatives of *Diaceratherium* now referable to *Brachydiceratherium* (see phylogenetic discussion), the most distinctive dental features of the Tagay rhinocerotid are a short premolar series (also observed in *Bd. shanwangense*), an enamel wrinkled and corrugated at the same time, crochets simple and lingual cingula usually absent and always reduced on P2-4, a protocone strongly constricted on M1, a lingual cingulum usually absent on lower premolars and always absent on lower molars, and the absence of d1/p1 at an adult stage (also observed in *Bd. shanwangense*). It can be further distinguished from *Bd. lamilloquense*, *Bd. lemanense*, *Bd. asphaltense*, and/or *Bd. aurelianense* by its I1s oval in occlusal view, the absence of labial cingulum on upper premolars and molars, the presence of a strong paracone fold on M1-2 and of a constricted hypocone on M1, and M3s with a triangular occlusal outline. Contrary to *Bd. lamilloquense*, the rhino from Tagay has a protoloph joined to the ectoloph on P2, but also a protocone and a hypocone lingually separate on P3-4 (molariform). With respect to *Bd. aurelianense*, it has no metaloph constriction on P2-4 and a protocone weakly developed on P2. In other words, dental remains from Tagay are strictly similar to those of *Bd. shanwangense* (Lu et al., 2021). They further have very close affinities with those of *Bd. aginense* and *Bd. intermedium* among *Brachydiceratherium* representatives. Nevertheless, the Tagay rhinocerotid differs from *Bd. aginense* in bearing a protocone and a hypocone lingually separate on P2 (molariform) and from both species in having a long metaloph on M1-2 and a posterior groove on M3. Even though postcranial elements are not known in all *Brachydiceratherium* species, the cervical vertebrae and/or limb bones from Tagay are perfectly matching those of *Bd. shanwangense* (Lu et al., 2021). They are also very similar to those of *Bd. aginense* and of *Bd. intermedium*, notably in terms of size and proportions. They differ, however, from all representatives of the genus (including the latter species), in having a scapular glenoid fossa with a straight medial border and a tibia-facet on the calcaneus, but no distal gutter on the humeral lateral epicondyle. It can be further distinguished from *Bd. lamilloquense*, *Bd. lemanense*, *Bd. asphaltense*, and/or *Bd. aurelianense* by a radius with a high posterior expansion of the scaphoid-facet, a femoral head hemispheric, an astragalus with a laterodistal expansion, the presence of very low-and-smooth intermediate reliefs on metapodials but also a long insertion of m. interossei on lateral metapodials. The Tagay rhinocerotid differs from *Bd. aginense* in bearing a shallow gutter for the m. extensor carpi on the radius, a posterior MtII-facet developed on the Mt3, but no contact between the cuboid and the MtIII, from *Bd. intermedium* in showing a right angle between the cuboid-facet and the base of the tuber calcanei on the calcaneus, and from both species in having a scaphoid with equal anterior and posterior heights, a short posterior tuberosity on the magnum, a wider astragalus (TD/H>1.2), and a fibula-facet on the calcaneus.

In fact, the Tagay rhinocerotid individual is identical in all aspects to corresponding specimens of the complete skeleton from the early Miocene of eastern China recently assigned to as *Diaceratherium shanwangense* by Lu et al. (2021). The only differences lie in the occipital crest being more concave in the Tagay skull than in the Shanwang one, following the description by Lu et al. (2021). This feature is likely to document either sexual dimorphism, ontogenetic variation, or interindividual variability. Accordingly, we consider unambiguously the Tagay rhinocerotid as documenting *Brachydiceratherium shanwangense*.

## PHYLOGENETIC ANALYSIS

We have first run a preliminary analysis (see supplementary materials: files S1 and S2), with 32 taxa i.e., the Tagay individual and *B. shanwangense* scored as two distinct terminals. In that analysis, these terminals differ in a single and only feature (char. 36: occipital crest concave in the former). Accordingly, we have merged them into a single terminal, for running the final analysis (see supplementary materials: files S3 and S4). A single most parsimonious tree is retrieved (length = 1316 steps; consistency index = 0.2698; retention index = 0.4918; Fig. 13; see supplementary files S3 and S4). Twenty-four characters are constant, due to their original definition for solving phylogenetic relationships within Elasmotheriina (Antoine, 2002), a rhinocerotid subtribe the representatives of which are not included here. Character distribution at each node and corresponding indices are detailed in the supplementary materials (file S4). Suprageneric relationships within Rhinocerotinae (i.e., the clade including Rhinocerotini + Aceratheriini) are consistent with those proposed by Antoine (2002, 2003), Antoine et al. (2010, 2022), Becker et al. (2013), Tissier et al. (2021), and Pandolfi et al. (2021): *Plesiaceratherium mirallesi* is the earliest offshoot among Rhinocerotinae (node 1; 26 unambiguous synapomorphies; Bremer Support [BS] > 5). Aceratheriini (node 3; nine unambiguous synapomorphies; BS = 2) and Rhinocerotini (node 5; eight unambiguous synapomorphies; BS = 2) are sister clades (node 2; 13 unambiguous synapomorphies; BS = 4). Rhinocerotina (node 6; 18 unambiguous synapomorphies; BS > 5) and Teleoceratina (node 13; five dental and postcranial unambiguous synapomorphies; BS = 1) are sister clades within Rhinocerotini (Fig. 13). Aceratheriini comprise *Alicornops simorrense* as a sister species to the (*Aceratherium incisivum*, *Acerorhinus zernowi*) clade (node 4). Rhinocerotina include the (*Lartetotherium sansaniense*, *Gaindatherium browni*) clade (node 7; seven unambiguous synapomorphies; BS = 5) as the first offshoot, then *Nesorhinus philippinensis* (node 8; seven unambiguous synapomorphies; BS = 3), and the living rhino species (node 9; nine unambiguous synapomorphies; BS = 2), with the *Rhinoceros* clade (node 10; four unambiguous synapomorphies; BS = 1) being sister group to the (*Dicerorhinus sumatrensis* plus African rhinos) clade (node 11; 13 unambiguous synapomorphies; BS = 3). The clade of living African rhinos is the most supported node of the tree (node 12; 38 unambiguous synapomorphies; BS > 5).

**Fig. 12.**
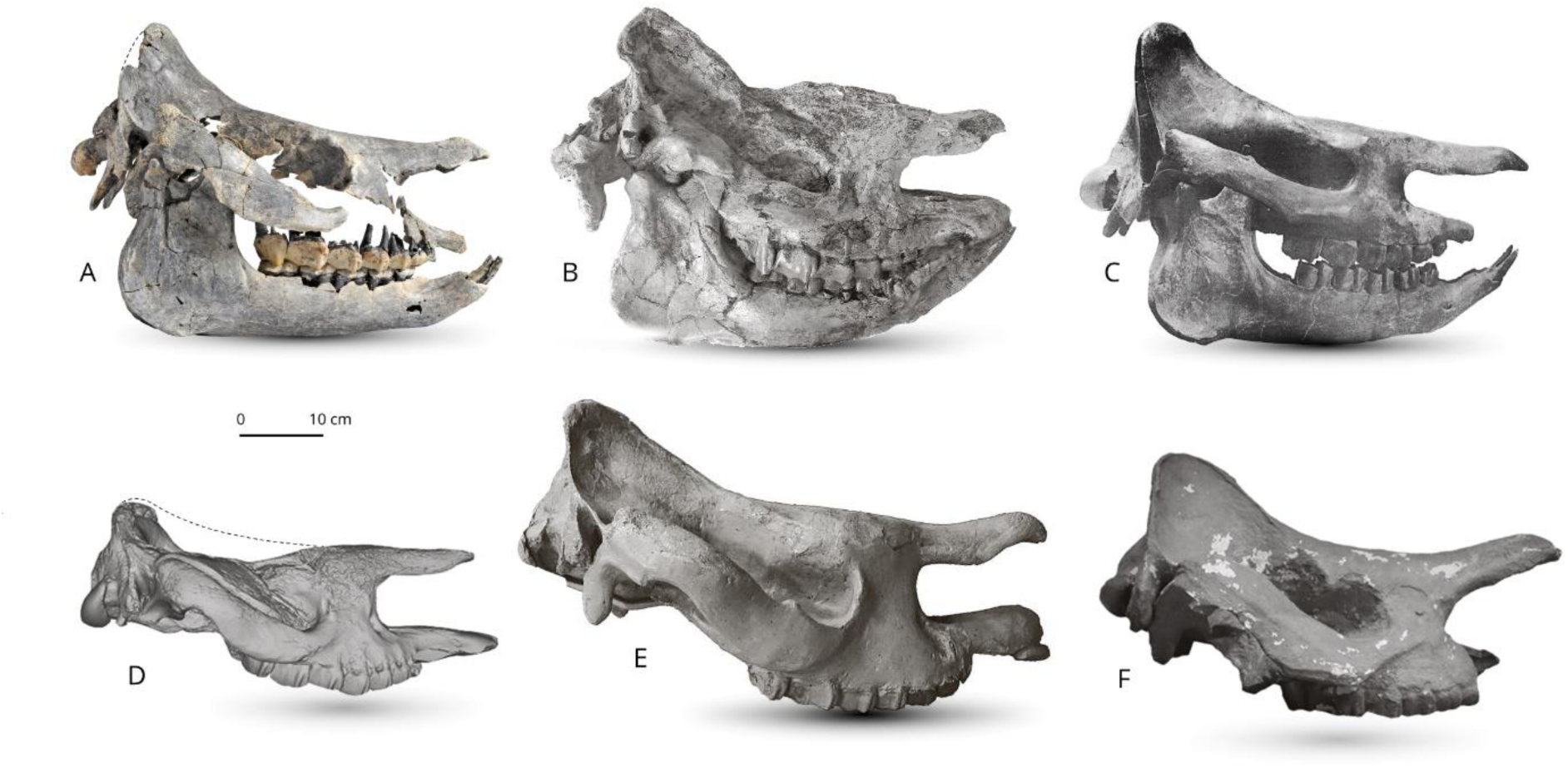
Skulls of different species of *Brachydiceratherium* in right lateral view. **A -***Brachydiceratherium shanwangense* from Tagay (Baikal Region, Russia, late early Miocene) №IZK79-1-08C-1/1; **B** -*Brachydiceratherium shanwangense* from Jijiazhuang locality STM 44–98 (deformed, mirrored) (MN4 - early Miocene, Shanwang Basin, Shandong Province, China) №MHNT.PAL.2013.0.1001; **C** - *Brachydiceratherium aginense* (Répelin, 1917) from Laugnac (MN2 - early Miocene, Lot-et-Garonne, France); **D** - *Brachydiceratherium lemanense* from Gannat (MN1 - early Miocene, France) №MNHN-AC-2375, holotype; **E** - *Brachydiceratherium asphaltense* (Depéret et Douxami, 1902) from Saulcet (MN1 - earliest Miocene, Allier, France). №NMB–Sau1662; **F** - *Brachydiceratherium aurelianense* from Neuville-aux-Bois (MN3 - early Miocene, France) №MHNT.PAL.2013.0.1001, cast of the holotype;

**Fig. 13.**
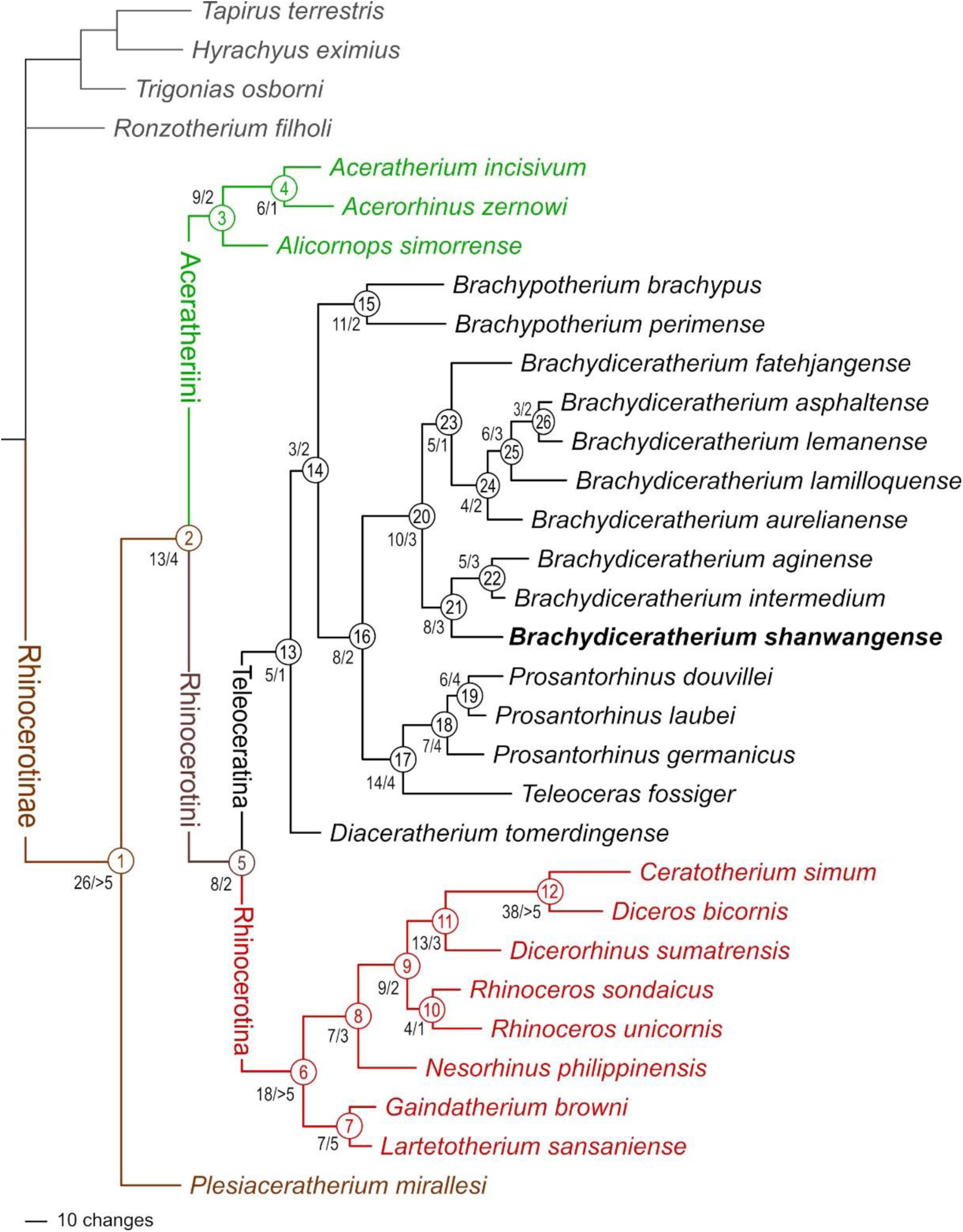
Phylogram of Rhinocerotinae, with a focus on Teleoceratina. Most parsimonious tree (1316 steps; consistency index = 0.2698; retention index = 0.4918), retrieved from 282 unweighted cranio-mandibular, dental, and postcranial characters scored in 31 tapirid and rhinocerotoid species (see S3 and S4). Node numbers appear in empty circles. Number of unambiguous synapomorphies/Bremer Support are indicated left to nodes.

In the next paragraphs, we will focus on the topology, node support (Bremer Support: BS), and apomorphy distribution regarding the Teleoceratina. The monophyly of the subtribe is weakly supported by five dental and postcranial unambiguous synapomorphies (BS = 1): I1 with an almond-shaped cross section, hypocone isolated by an anterior constriction on M2, ulna with the olecranon and the diaphysis forming a closed angle, robust limbs, and lateral metapodials with insertions of the m. interossei short. The earliest-diverging teleoceratine is *Diaceratherium tomerdingense*. This species is defined by ten dental and postcranial autapomorphies (teeth with enamel wrinkled and roots separate, P2-3 with an antecrochet usually absent, M1-2 with a metaloph short, M2 with a mesostyle, humerus without a distal gutter on the lateral epicondyle, semilunate with a distal border of the anterior side rounded, trapezoid with a proximal border asymmetric in anterior view, unciform with a posterior expansion of the pyramidal-facet always present, and trapezium-facet always absent on the McII; Table 6). Next node (node 14) segregates the *Brachypotherium* clade (node 15) from all other teleoceratines scored here (node 16). Node 14 (BS = 2) is weakly supported by three postcranial unambiguous synapomorphies (proximal ulna-radius facets usually fused, gutter for the m. extensor carpi weakly developed on the radius, and McII with anterior and posterior McIII-facets fused). Eleven cranio-mandibular, dental, and postcranial synapomorphies define *Brachypotherium* (node 15; BS = 2): occipital condyle without a median ridge, mandibular symphysis very massive, labial cingulum usually present on upper premolars and always present on upper molars, lower cheek teeth with a flat ectolophid, lower premolars with a lingual opening of the posterior valley U-shaped, p2 with a paraconid reduced, radius-ulna with a second distal articulation, pyramidal with a distal semilunate-facet asymmetric, posterior facet always absent on the McII-McIII, and fibula-facet oblique on the astragalus. The Bremer Support is low, due to an alternative topology with *B. perimense* being sister taxon to the (*B. brachypus*, node 16) clade appearing at 1317 steps. *Brachypotherium brachypus* is particularly well differentiated, with 27 unambiguous cranio-mandibular, dental, and postcranial autapomorphies each (see Table 6). From node 16 diverge two clades, with (*Teleoceras* plus *Prosantorhinus*) on the one hand (node 17), and all species classically assigned to *Diaceratherium* except the type species (node 20). Node 16 (BS = 2) is supported by eight cranio-dental and postcranial unambiguous synapomorphies: vomer rounded, protocone constriction usually absent on P3-4, antecrochet always present on P4, lingual cingulum always present on lower premolars, pyramidal- and McV-facets always separate on the unciform, McIV with a trapezoid outline in proximal view, calcaneus-facets 2 and 3 always independent on the astragalus, and fibula-facet always present on the calcaneus. Node 17 (BS = 4) places the highly-divergent *Teleoceras fossiger* (39 cranio-mandibular, dental, and postcranial unambiguous autapomorphies; Table 6) as sister species to *Prosantorhinus*, through 14 cranio-mandibular, dental, and postcranial synapomorphies: base of the processus zygomaticus maxillary low on the maxilla, zygomatic arch high, articular tubercle of the squamosal concave, lingual groove (sulcus mylohyoideus) absent on the corpus mandibulae, metaloph transverse and protoloph sometimes interrupted on P2, mesostyle present on M2, d2 with a posterior valley usually open, scapula spatulated and with a medial border straight on the glenoid fossa, a trochanter major low on the femur, MtII-facet always absent and cuboid-facet present on the MtIII, and metapodials with high and acute intermediate reliefs. *Prosantorhinus* (node 18; BS = 4) is monophyletic, with *P. germanicus* (thirteen cranio-dental unambiguous autapomorphies; Table 6) as the first offshoot (node 18) and *P. laubei* and *P. douvillei* being sister species (node 19). The monophyly of *Prosantorhinus* is supported by seven cranio-dental unambiguous synapomorphies, some being optimised in *P. laubei* (no cranial remains available; Heissig & Fejfar, 2007): lateral apophysis present on the nasals, median nasal horn present (probably in males), presence of a sagittal fronto-parietal crest, of a posterior groove on the processus zygomaticus of the squamosal, of a metacone fold on M1-2, of an unconstricted metaloph on M1, and of an ectolophid fold on d2-3. *Prosantorhinus douvillei* (nine unambiguous dental autapomorphies; Table 6) and *P. laubei* (six unambiguous dental autapomorphies; Table 6) share six dental and postcranial unambiguous synapomorphies (node 19; BS = 4): protocone unconstricted on P3-4 and M3, metaloph unconstricted on M2, labial cingulum always present on lower molars, lingual groove always present on d3, and expansion of the calcaneus-facet 1 always high and narrow on the astragalus.

**Table 6.**
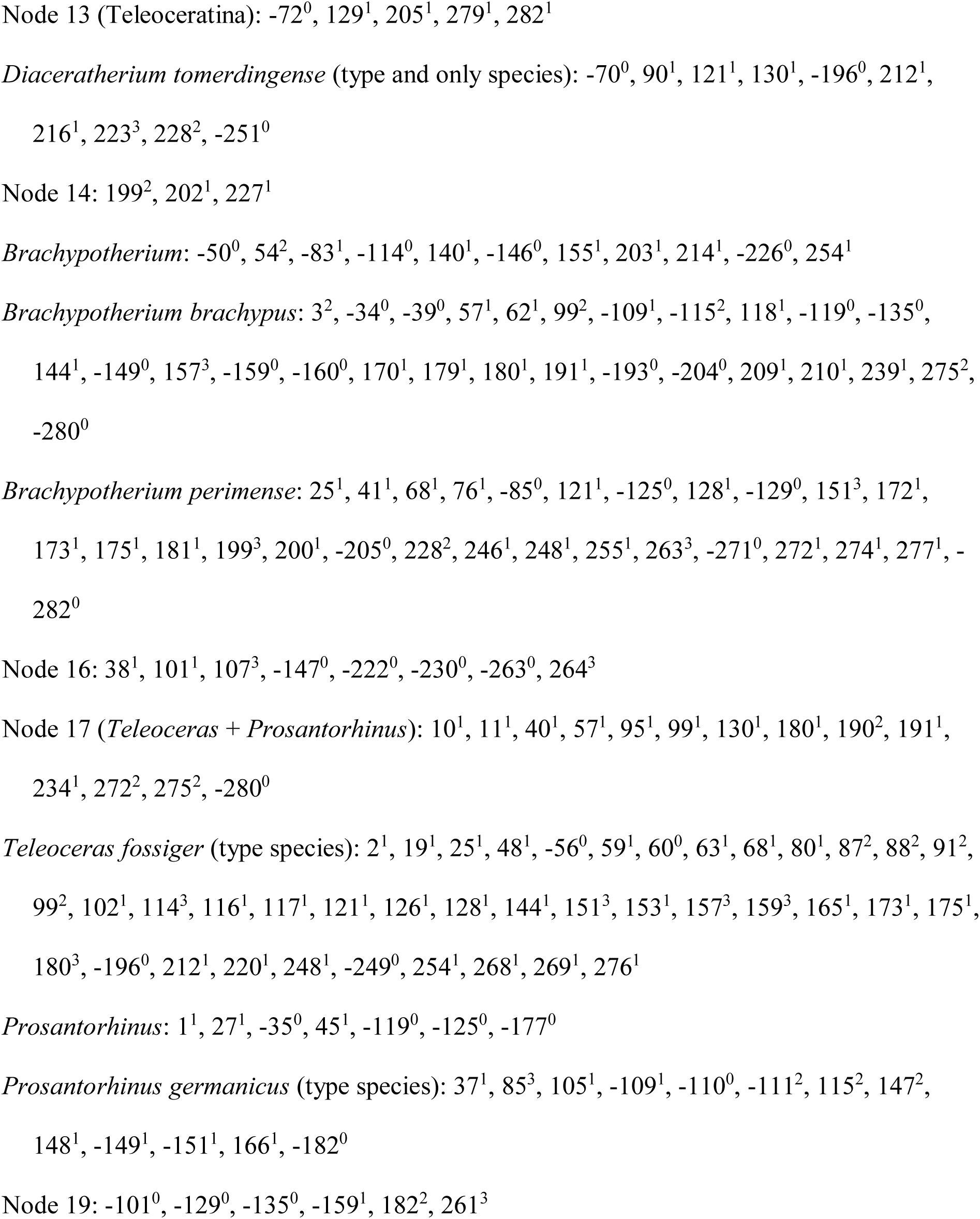
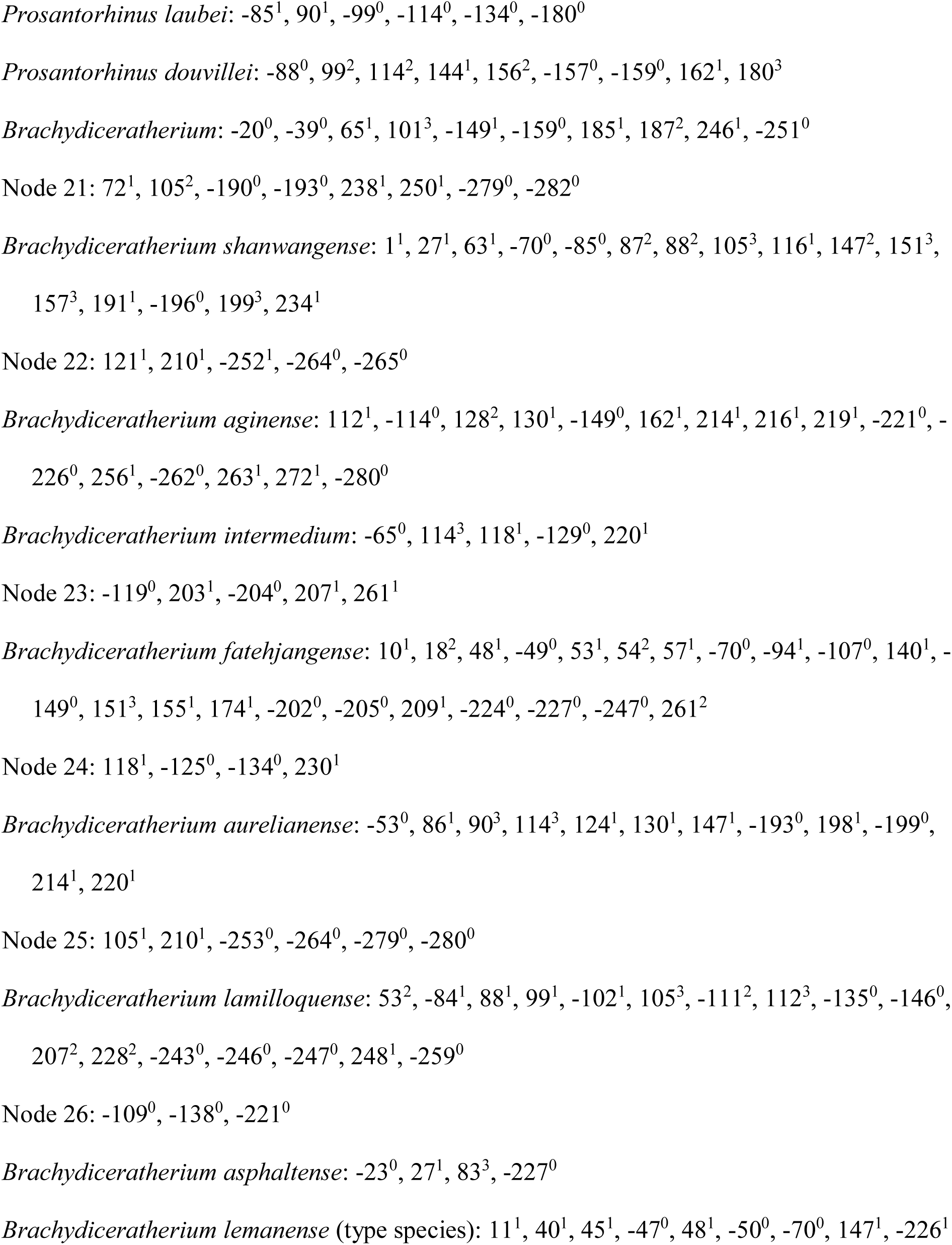
Distribution of unambiguous apomorphic characters (synapomorphies and autapomorphies, including reversals) among teleoceratine rhinocerotids, as retrieved in the current phylogenetic analysis. Node numbers match those of Fig. 13. Binominal combinations are as detailed in the Discussion.

Node 20 (BS = 3) gathers eight terminal taxa (Fig. 13). It is supported by ten cranio-dental and postcranial synapomorphies: nuchal tubercle small, articular tubercle smooth on the squamosal, cement present on cheek teeth, protocone always constricted on P3-4, labial cingulum usually absent on lower premolars and always present on lower molars, foramen vertebrale lateralis present and axis-facets transversally concave on the atlas, postero-distal apophysis low on the tibia, and latero-distal gutter located posteriorly on the fibula. Two clades diverge from node 20. The first one (node 21, BS = 3) gathers *Brachydiceratherium shanwangense*, *Bd*. *aginense*, and *Bd. intermedium*, based on eight dental and postcranial synapomorphies: I1 with an oval occlusal outline, labial cingulum always absent on upper premolars, crista usually present on P3, scapula elongated, fossa olecrani high on the humerus, fovea capitis low and wide on the femur, latero-distal gutter deep on the fibula, limbs slender, and insertions for the m. interossei long on lateral metapodials. Most of them are optimised in *Bd. intermedium*. *Brachydiceratherium shanwangense* is well diagnosed, with sixteen cranio-dental and postcranial unambiguous synapomorphies: lateral apophysis present on the nasals, median nasal horn present on the nasals, premolar series short with respect to the molar series, roots distinct on the cheek teeth, crochet always simple and lingual cingulum usually absent and always reduced on P2-4, crista always present on P3, protocone strongly constricted on M1-2, lingual cingulum usually absent on lower premolars and always absent on lower molars, d1/p1 absent in adults, glenoid fossa with a medial border straight on the scapula, distal gutter absent on the lateral epicondyle of the humerus, proximal radius-ulna facets always fused, and trochanter major low on the femur. Node 22 (BS = 3) is supported by five dental and postcranial unambiguous synapomorphies: metaloph short on M1-2, posterior height exceeding the anterior height on the scaphoid, astragalus almost as high as wide (TD/H ratio between 1 and 1.2), and tibia- and fibula-facets absent on the calcaneus.

*Brachydiceratherium intermedium* (five dental and postcranial unambiguous autapomorphies; Table 6) is less derived than *Bd. aginense* (16 dental and postcranial unambiguous autapomorphies; Table 6), which probably reflects the strong contrast in the completeness of their hypodigms (e.g., no indisputable cranial remains are documented for *Bd. intermedium*).

The second clade diverging from the node 20 (i.e., node 23) places *Bd. fatehjangense* as a sister taxon to (*Bd. aurelianense*, (*Bd. lamilloquense*, (*Bd. lemanense*, *Bd. asphaltense*))). All the corresponding nodes are weakly supported (1 ≤ BS ≤3), with low numbers of unambiguous synapomorphies (ranging from three to six). Node 23 is the least-supported one (BS = 1), with five dental and postcranial synapomorphies (metacone fold present on M1-2, second distal radius-ulna articulation present, posterior expansion of the scaphoid-facet low on the radius, postero-proximal semilunate-facet usually absent on the scaphoid, and expansion of the calcaneus-facet 1 usually wide and low on the astragalus).

## DISCUSSION

### Ontogenetic age and sex

Both the complete dental eruption and the wear stages of upper and lower teeth concur to consider this individual as an adult, most likely ∼7-15 years old (with reference to recent rhinos; e.g., Hillman-Smith et al., 1986; Hullot et al., 2020). In the absence of I1s (usually highly dimorphic in teleoceratines; see Antoine, 2002 regarding *Prosantorhinus douvillei*), and due to the fragmentary state of i2s, it is not possible to determine its sex.

### Taxonomic inferences

Surprisingly, *Diaceratherium tomerdingense* Dietrich, 1931 is retrieved as the first offshoot among Teleoceratina (Fig. 13). Moreover, the assignment of this hornless and robust-limbed rhinocerotine to the subtribe is not well supported at all (BS = 1): in other words, this species could be closely related to Rhinocerotina instead among Rhinocerotini, as suggested by some of its peculiar features, retrieved as autapomorphies in the current analysis (metaloph short on M1-2; distal gutter on the lateral epicondyle absent on the humerus, distal border of the anterior side of the semilunate rounded, and trapezium-facet absent on the McII). Accordingly, and taking into account both the topology of the most parsimonious tree and the character distribution along its branches, we propose that *Diaceratherium* Dietrich, 1931 shall be restricted to the type species.

Indeed, all other species previously assigned to *Diaceratherium* in the last decades form a well-supported clade remote from the type species (Fig. 13). This clade is split into two sister clades encompassing three and five species, respectively (*D. shanwangense*, *D. aginense*, and *D. intermedium*; *D. fatehjangense*, *D. aurelianense*, *D. lamilloquense*, *D. asphaltense*, and *D. lemanense*). Except for *D. lamilloquense* Michel, 1987, these species were originally or subsequently assigned to pre-existing genera, i) either unambiguously non-related to Teleoceratina, such as *Aceratherium* (*D. lemanense*), *Diceratherium* (*D. asphaltense*, *D. lemanense*), *Aprotodon* (*D. fatehjangense*), *Chilotherium* or *Subchilotherium* (*D. intermedium*), and *Plesiaceratherium* (*D. shanwangense*), or ii) among Teleoceratina, with *Teleoceras* and/or *Brachypotherium* (*D. aginense*, *D. aurelianense*, *D. shanwangense*, and *D. fatehjangense*). Finally, and to our knowledge, the only species belonging to this clade for which a genus-group name has been unambiguously proposed is *D. lemanense*. Indeed, Lavocat (1951) has erected the subgenus *Brachydiceratherium* for “*Acerotherium lemanense* Pomel, 1853”. Interestingly, Lavocat did assign these species and subgenus to *Diceratherium* Marsh, 1875, a genus consistently assigned to Elasmotheriinae in the last decades (e.g., Antoine, 2002). We propose that all these eight species be assigned to *Brachydiceratherium* Lavocat, 1951, especially as the five-species clade, with *D. fatehjangense*, *D. aurelianense*, *D. lamilloquense*, *D. asphaltense*, and *D. lemanense*, is not well supported (BS = 1; 5 unambiguous synapomorphies). Noteworthily, *D. asphaltense* and *D. lemanense* are sister species in the most parsimonious tree, with a low number of morpho-anatomical discrepancies. It should be noted that Jame et al. (2019) consider both species as being distinct, based on a wide array of cranio-dental and postcranial features.

Other teleoceratine genera are monophyletic in the present analysis. *Brachypotherium* Roger, 1904 includes *B. brachypus* and *B. perimense* and this genus is a sister group to a clade gathering *Teleoceras* Hatcher, 1894 plus *Prosantorhinus* Heissig, 1974 on one branch and *Brachydiceratherium* on the other one (see above).

### Historical biogeography of Eurasian teleoceratines

During early Miocene times, Teleoceratina were particularly species-rich in Eurasia, with 5–8 coeval species in any time slices (Fig. 14). A common thread between *Brachypotherium*, *Brachydiceratherium*, and *Prosantorhinus* is their huge geographical range at the generic level, encompassing most of the Eurasian landmasses for the latter two genera (e.g., Heissig, 1999; Antoine et al., 2010, 2013), plus Afro-Arabia for *Brachypotherium* (e.g., Hooijer, 1963, Geraads & Miller, 2013; Pandolfi & Rook, 2019). An early representative of *Brachydiceratherium* has been recognised in Thailand (*Bd*. cf. *lamilloquense*; Marivaux et al., 2004). It has the closest affinities with *Bd. lamilloquense*, from the late Oligocene of Western Europe (Fig. 15). To our knowledge, no occurrence has been reported between both areas for this species. *Prosantorhinus* has a similar geographical range, extending from Western Europe (*P. germanicus* and *P. douvillei*; Heissig, 1972b; Antoine et al., 2000; Heissig, 2017) and Central Europe (*P. laubei*; Heissig & Fejfar, 2007) to Southern Pakistan (*P. shahbazi*; Antoine et al., 2010, 2013). If confirmed, the recognition of *Bd. fatehjangense* in lower Miocene beds of the Turgai region in Kazakhstan, previously described as a representative of *Bd. aurelianense* by Borissiak (1927) and Lu et al. (2021), would considerably expand latitudinally the range of this species, previously restricted to the Indian Subcontinent. It would then be documented on both sides of the Himalayas (Fig. 15). The ubiquitous distributions of most teleoceratine taxa likely underline ultra-generalist ecological preferences (Hullot et al., 2021). Moreover, such ranges seemingly support the absence of efficient ecological and geographical barriers at the Eurasian scale for the concerned teleoceratines, at least by early Miocene times (Fig. 15).

**Fig. 14.**
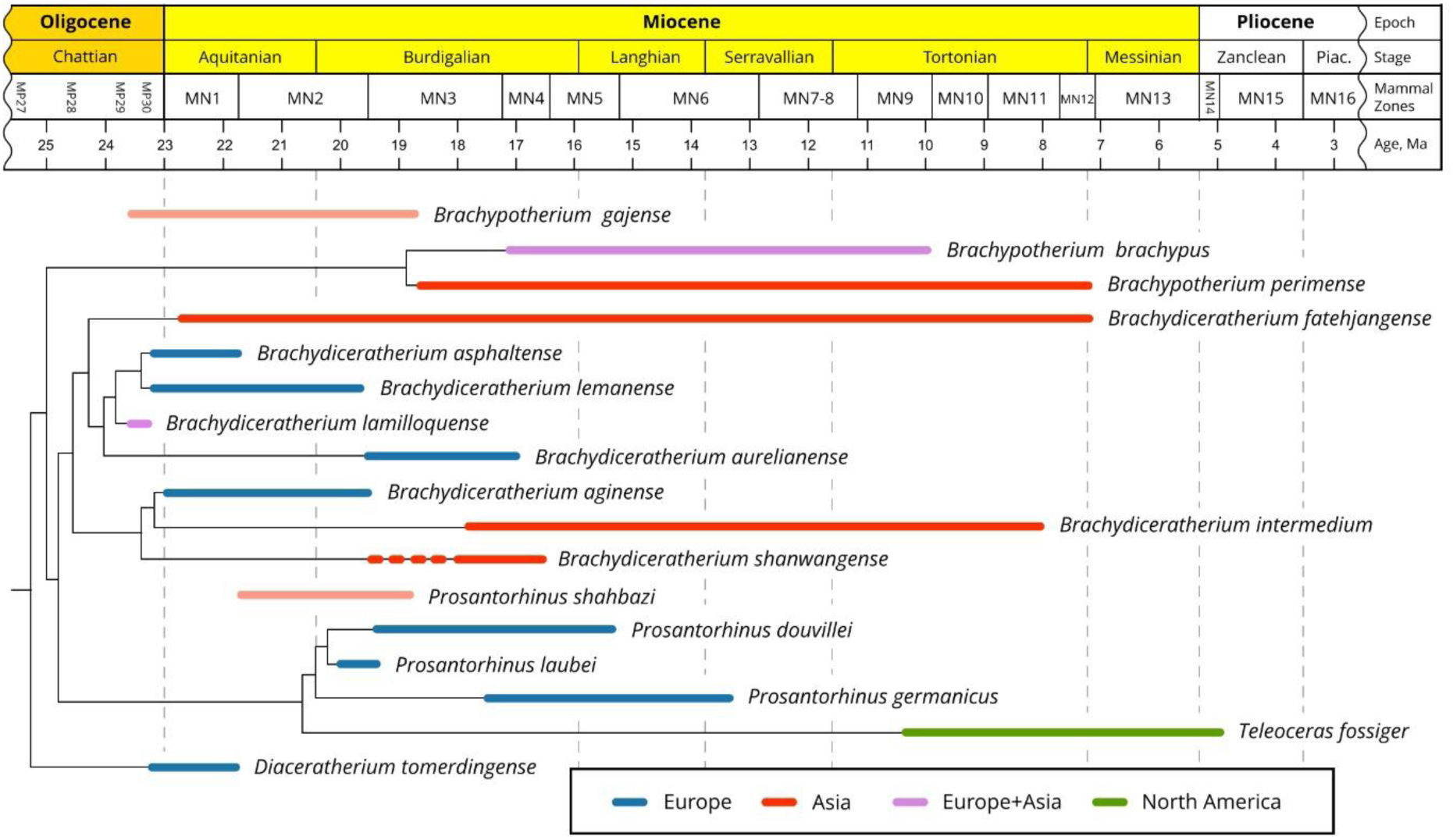
Phylogenetic relationships of Teleoceratina versus time (see Fig. 13), with new combinations. Although they were not included in the current parsimony analysis, the temporal distributions of *Brachypotherium gajense* and *Prosantorhinus shahbazi* are provided here, as these species might bridge a stratigraphic gap for the concerned genera. Red dotted line for *B. shanwangense* stands for the age uncertainty of Tagay locality (MN3-5)

**Fig. 15.**
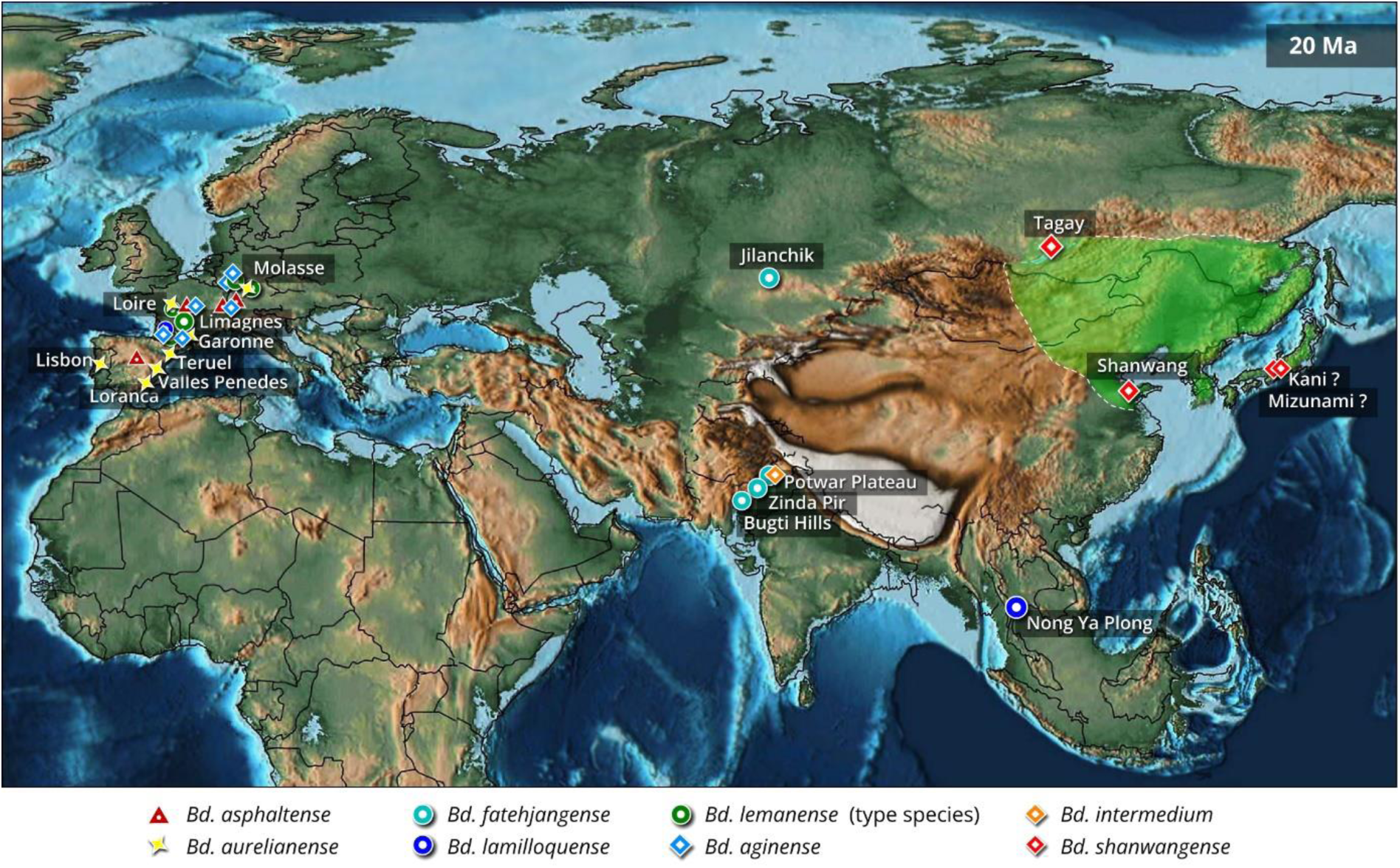
Paleomap of Eurasia by early Miocene times (∼20 Ma), showing the main occurrences of representatives of the teleoceratine rhinocerotid *Brachydiceratherium*, at the basin scale (apart from Tagay, Shanwang, and Nong Ya Plong localities). The Green area depicts the interpolated geographical range of *B. shanwangense* (with possible occurrences on Honshu Island, Japan). Based on data from Borissiak (1927), Cerdeño (1993), Antoine et al. (2000, 2013), Becker et al. (2009), Antoine & Becker (2013), Tomida et al. (2013), Jame et al. (2019), Handa (2020), Lu et al. (2021), Antoine (in press), and the present work. PalaeoAtlas by Scotese (2016, under cc 4.0 license) with added paleomap for the Baikal area (Mats et al., 2011).

Moreover, ghost lineages within *Brachypotherium* and *Prosantorhinus* (Fig. 14) are likely to be bridged by *B. gajense* and *P. shahbazi*, from the latest Oligocene–earliest Miocene and the early Miocene of Pakistan, respectively (for further discussion, see Antoine et al., 2013 and Antoine, in press).

*Brachydiceratherium shanwangense* was previously only documented at Shanwang, eastern China (N32°, E116.5°). The well-supported specific assignment of the Tagay rhinoceros (N53°, E107.5°) points to an unsuspectedly wide geographical range for this species, further pleading for both a low climatic and environmental gradient in the concerned area at that time and very broad ecological preferences for this species (Fig. 15). Moreover, it can be suspected that the smallest teleoceratine remains described over the early Miocene interval in Japan (Kani and Mizunami formations) and referred to the *Brachypotherium pugnator* (Matsumoto, 1921), otherwise of gigantic dimensions (Fukuchi & Kawai, 2011; Tomida et al., 2013; Handa, 2020), may have particularly close affinities with those of *Bd. shanwangense*. More generally, the concerned Japanese assemblages are very similar to the Tagay and Shanwang ones (e.g., with the equid *Anchitherium* cf. *gobiense*, the proboscidean *Gomphotherium annectens*, and the beaver *Youngofiber sinensis*; Qiu & Qiu, 2013), thus strengthening the existence of a single eastern Asian biogeographical province at mid latitudes at that time (Fig. 15). Indeed, closed forest environments under a subtropical climate, with precipitation averaging ca. 1500 mm per year, are reported for the Shanwang Basin based on early Miocene floras and vertebrates (Lu et al., 2021). The same proxies allow for considering the Tagay area as a lake, also surrounded by dense forests under subtropical conditions, with precipitation averaging ca. 1000-1500 mm per year (Logachev et al, 1964; Belova, 1985; Sizov & Klementiev, 2015).

## CONCLUSIONS

The numerous associated features documented and scored in the Tagay rhinocerotid skeleton have allowed for assigning it to the same teleoceratine species (*Brachydiceratherium shanwangense*) as in Shanwang, eastern China. These remains further contribute to a refined depiction of phylogenetic relationships and to a revision of generic assignments among Eurasian Teleoceratina.

The genus *Diaceratherium* Dietrich, 1931 should be restricted to the type species, *Diaceratherium tomerdingense* Dietrich, 1931. This monotypic genus is the first offshoot within Teleoceratina. Our results support the reappraisal of *Brachydiceratherium* Lavocat, 1951, with eight assigned species: *Brachydiceratherium lemanense* (Pomel, 1853), *Brachydiceratherium aurelianense* (Nouel, 1866), *Brachydiceratherium intermedium* (Lydekker, 1884), *Brachydiceratherium asphaltense* (Depéret & Douxami, 1902), *Brachydiceratherium fatehjangense* (Pilgrim, 1910), *Brachydiceratherium aginense* (Répelin, 1917), *Brachydiceratherium shanwangense* (Wang, 1965) and *Brachydiceratherium lamilloquense* Michel, 1983. *Brachydiceratherium* is a sister group to a clade encompassing *Prosantorhinus* and the North American genus *Teleoceras*. *Brachypotherium* is more closely related to the latter three genera than to *Diaceratherium*.

All Old World teleoceratines have extended geographical distributions at the genus level, which is also true for some species, such as the late Oligocene *Brachydiceratherium lamilloquense* and the early Miocene *Brachydiceratherium shanwangense*. The latter range supports the existence of a single eastern Asian biogeographical province at mid latitudes at that time for such megaherbivores.

## Supporting information

Character matrix for the preliminary phylogenetic analysis

Output log text of the preliminary phylogenetic analysis

Character matrix for the final phylogenetic analysis

Output log text of the final phylogenetic analysis

Measurements for Brachydiceratherium shanwangense from Tagay site. XLSX

Measurements for Brachydiceratherium shanwangense from Tagay site. PDF

## ACKNOWLEDGMENTS

We are grateful to our colleagues for having participated in the 2008-2021 excavations. Special thanks to Gennady Turkin for logistical support and for assistance in the field. Valeria Burova and Ekaterina Nikulina are acknowledged for working on the skeleton reconstruction. Géraldine Véron and Christine Argot (Muséum National d’Histoire Naturelle (Paris, France) kindly provided access to zoological and paleontological collections under their care. We deeply acknowledge the Recommender, Faysal Bibi, and all three reviewers (Jérémy Tissier, Deng Tao, and Panagiotis Kampouridis), for their thorough revisions and constructive remarks on a widely-improvable previous version of the manuscript.

## SUPPORTING INFORMATION

Additional supporting information may be found online both in the supporting information tab for this article and on Morphobank for matrices (http://morphobank.org/permalink/?P5029). Supplementary Files:

- **S1**. Character matrix for the preliminary phylogenetic analysis, including 282 cranial, dental, and postcranial characters controlled on 32 terminal taxa (one tapirid, rhinocerotoids, and rhinocerotids), with Tagay rhinoceros and “*Diaceratherium shanwangense*” as separate terminals.
- **S2.** Output log text of the preliminary phylogenetic analysis (282 characters and 32 taxa).
- **S3**. Character matrix for the final phylogenetic analysis, including 282 cranial, dental, and postcranial characters controlled on 31 terminal taxa (one tapirid, rhinocerotoids, and rhinocerotids).
- **S4.** Output log text of the final phylogenetic analysis, with Bremer Support
- **S5** Measurements for *Brachydiceratherium shanwangense* from Tagay site.

