## Supplementary material for "An Early Miocene skeleton of *Brachydiceratherium* Lavocat, 1951 (Mammalia, Perissodactyla) from the Baikal area, Russia, and a revised phylogeny of Eurasian teleoceratines": Measurements for Brachydiceratherium shanwangense from Tagay site. PDF

**SKULL**

| 1 | 2 | 3 | 4 | 5 | 6 | 7 | 8 | 9 | 10 | 11 | 12 | 13 | 14 | 15 | 16 | 17 | 18 | 19 | 20 |
|---|---|---|---|---|---|---|---|---|---|---|---|---|---|---|---|---|---|---|---|
| 505 | 540 | 455.8 | 174 | 125 | 249.6 | 272 | 293.6 | 72.6 | - | - | - | 254 | 219 | 160 | 224.8 | 26.7 | 179 | 189 | - |

| 21 | 22 | 23 | 24 | 25 | 26 | 27 | 28 | 29 | 30 | 31 | 32 |
|---|---|---|---|---|---|---|---|---|---|---|---|
| 308.2 | 69.7 | 144.6 | - | 140 | 155 | 156 | - | - | - | 43 | 112.4 |

**Cranial measurements of Brachydiceratherium shanwangense, from Tagay, Early Miocene of Eastern Siberia, in mm.**

Numbers coincide with measurements as defined and illustrated by Guérin (1980, fig. 1, table 1).

Guérin C. 1980. Les rhinocéros (Mammalia, Perissodactyla) du Miocène terminal au Pléistocène supérieur en Europe occidentale. Comparaison avec les espèces actuelles. Documents des Laboratoires de Géologie de Lyon 79: 1–1185.

**MANDIBLE**

| 1 | 2 | 3 | 4 | 5 | 6 | 7 | 8 | 9 | 10 | 11 | 12 | 13 | 14 | 15 | 16 | L. diast. | D I2 |
|---|---|---|---|---|---|---|---|---|---|---|---|---|---|---|---|---|---|
| 467.4 | 136.8 | 68 | 68 | 70.7 | 75 | 76.4 | 77.7 | 38.1 | 41.3 | 118.2 | - | 136.8 | 96.2 | 203.3 | 229.2 | 51 | - |

**DENTAL MEASUREMENTS**

| | | $P_1$ | $P_2$ | $P_3$ | $P_4$ | $M_1$ | $M_2$ | $M_3$ | $p_2$ | $p_3$ | $p_4$ | $m_1$ | $m_2$ | $m_3$ |
|---|---|---|---|---|---|---|---|---|---|---|---|---|---|---|
| Left | L | - | 25.1 | 31.5 | 35.3 | 47.0 | 50.5 | - | - | 28.0 | 31.8 | 38.6 | 43.8 | 43.9 |
| | W | - | 31.5 | 41.1 | 47.7 | 51.7 | 54.2 | - | - | 21.8 | 25.2 | 27.6 | 29.8 | 30.5 |
| | H | - | 17.1 | 21.7 | 28.3 | 30.5 | 37.5 | - | - | 18.2 | 23.2 | 28.8 | 30.2 | 29.8 |
| Right | L | 19.3 | 27.2 | 29.0 | 35.5 | 47.1 | 50.6 | 42.1 | - | 29.8 | 32.0 | 37.7 | 44.1 | 45.4 |
| | W | 14.5 | 31.9 | 40.0 | 47.6 | 51.4 | 56.2 | 46.1 | - | 21.9 | 27.2 | 28.1 | 30.0 | 30.3 |
| | H | 8.8 | 18.0 | 21.0 | 27.0 | 29.7 | 37.6 | 37.8 | - | 19.8 | 23.0 | 24.8 | 28.7 | 27.0 |

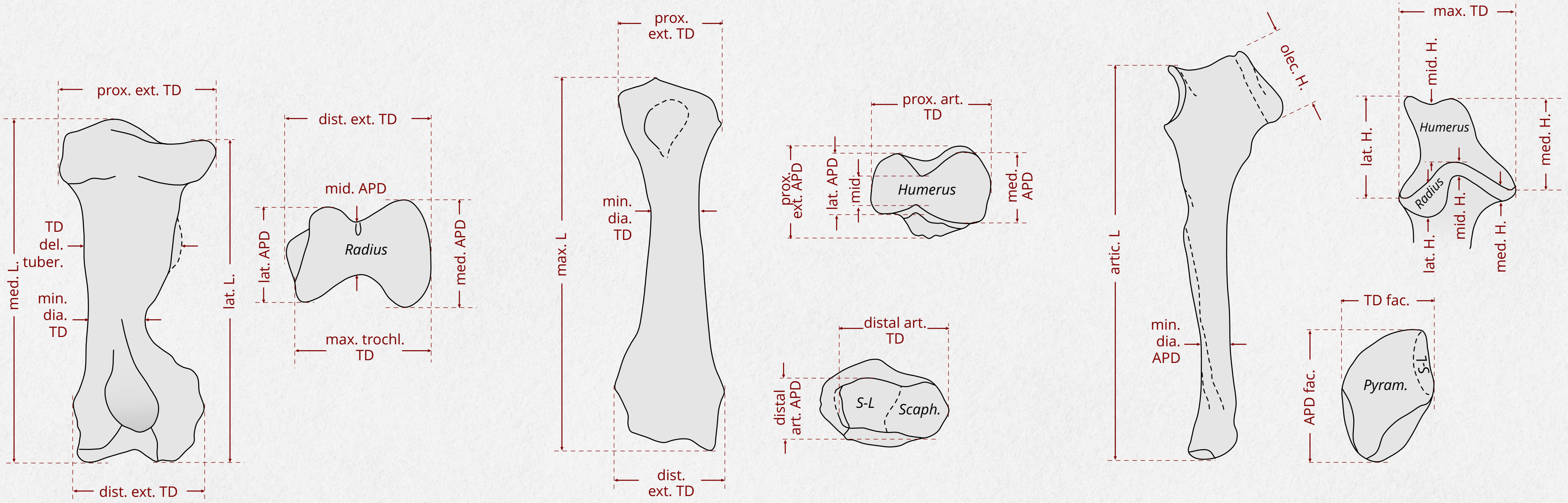


**HUMERUS** **RADIUS** **ULNA**

| HUMERUS | L | | proximal extremity | | TD del. tuber. | diaphysis | | distale extremity | | TD trochlea | | | APD trochlea | | |
|---|---|---|---|---|---|---|---|---|---|---|---|---|---|---|---|
| | med. | lat. | TD | APD | | min. TD | APD | TD | APD | max. | med. | lat. | med. | mid. | lat. |
| | 393 | 400.3 | 120.7 | 126.8 | 104 | 54.1 | 55.1 | 127.6 | 99 | 92.5 | 89.4 | 65 | 75 | 42 | 57.2 |

| RADIUS | max. L | proximal extremity | | proximal articulation | | | | diaphysis | | distale extremity | | distale articulation | |
|---|---|---|---|---|---|---|---|---|---|---|---|---|---|
| | | TD | APD | TD | med. APD | mid. APD | lat. APD | min. TD | min. APD | TD | APD | TD | APD |
| | 324.1 | 88.1 | 58 | 87 | 47.7 | 40.3 | 35.7 | 39 | 36.5 | 95.5 | 54.4 | 84.6 | 40 |

| ULNA | artic. L | olecranon | | humeral cochlea | | | | Radius fac. | | | diaphysis | | Pyram. fac. | | S.-L. fac. | | dist. Radius fac. | |
|---|---|---|---|---|---|---|---|---|---|---|---|---|---|---|---|---|---|---|
| | | TD | H | max. TD | med. H. | mid. H. | lat. H. | med. H. | mid. H. | lat. H. | TD | APD | TD | APD | TD | APD | APD | H |
| | 353.9 | 67.5 | 79.8 | 81.3 | 57.6 | 44.7 | 66.5 | 10.5 | 9.8 | 24.9 | 46.3 | 31.4 | 32.1 | 46.5 | not available | | 16.5 | 7.2 |

### THE CARPAL BONES
MEASUREMENTS

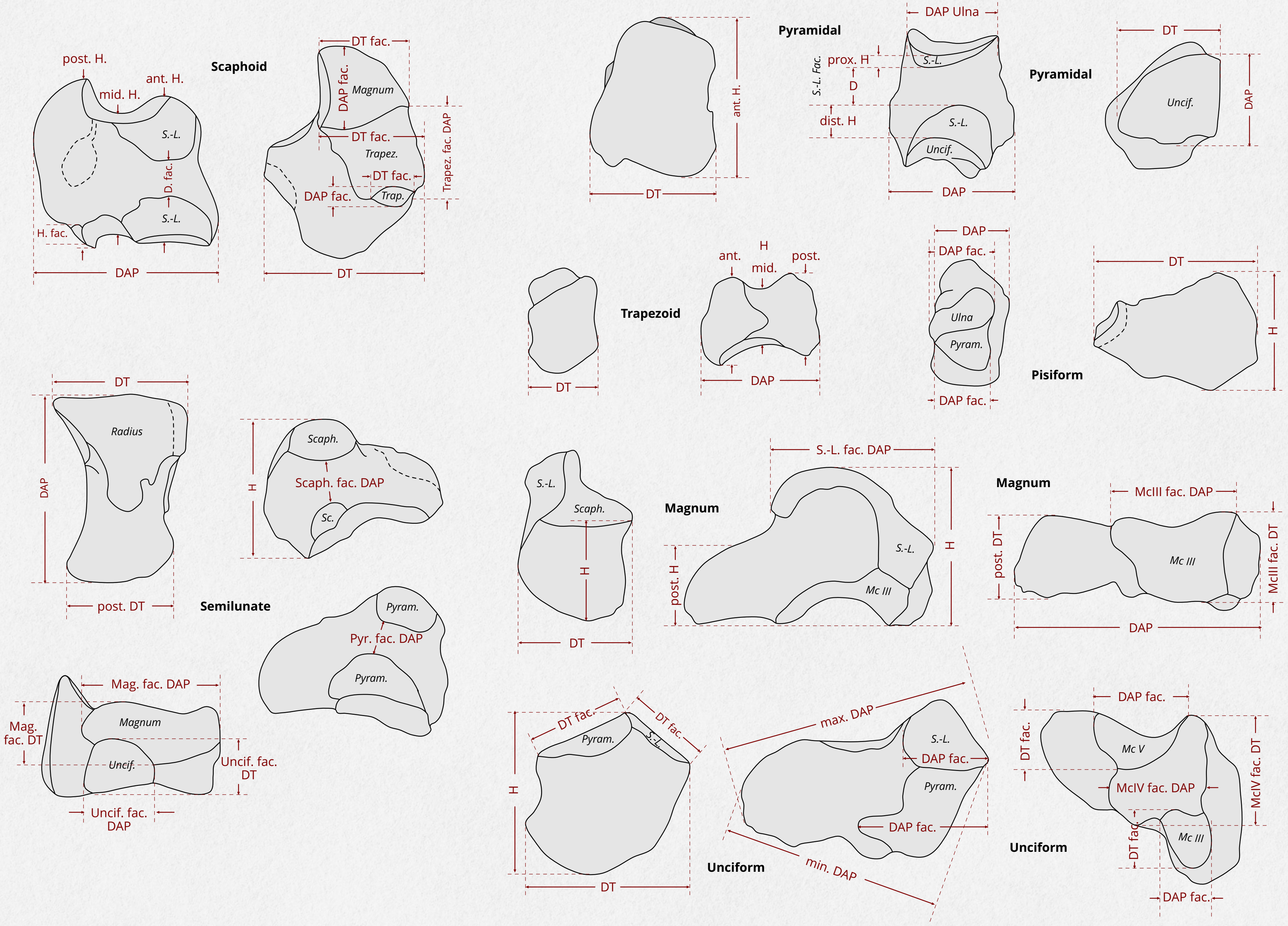

**Scaphoid**

| TD | APD | ant. H | mid. H | post. H | Radius fac. APD | Trapezium fac. | | Trapezoid fac. | | Magnum fac. | | S.-L. fac. D |
|---|---|---|---|---|---|---|---|---|---|---|---|---|
| | | | | | | TD | APD | TD | APD | TD | APD | |
| 52.7 | 73.6 | 49 | 34.4 | 46.4 | 39 | 18.6 | 12.4 | 29.1 | 39.7 | 30.4 | 31.3 | 11.1 |

**Semilunate**

| TD | APD | H | post. TD | Magnum fac. | | Unciform fac. | | Scaph. fac. D | Pyram. fac. D |
|---|---|---|---|---|---|---|---|---|---|
| | | | | TD | APD | TD | APD | | |
| 36.1 | 63.6 | 47.1 | 28.1 | 22.8 | 47.7 | 17.8 | 33 | 10.7 | 6.3 |

**Pyramidal**

| TD | APD | ant. H | Ulna fac. APD | Unciform fac. | | Semilunate fac. | | |
|---|---|---|---|---|---|---|---|---|
| | | | | TD | APD | D | prox. H | dist. H |
| 45.3 | 44 | 56.5 | 32.6 | 38.9 | 28.5 | 6.8 | 12.7 | 9.5 |

**Trapezoid**

| TD | APD | Height | | | Trap. fac. APD min. |
|---|---|---|---|---|---|
| | | ant. | mid. | post. | |
| 34.4 | 35 | 27.6 | 19.6 | 29.4 | 20 |

**Magnum**

| TD | ant. H | H | APD | S.-L. fac. APD | Mc III fac. | | post. tuber. | |
|---|---|---|---|---|---|---|---|---|
| | | | | | TD | APD | TD | H |
| 49 | 26.1 | 47.7 | 74.1 | 40.2 | 43.5 | 42.2 | 17.6 | 25.9 |

**Pisiform**

| TD | APD | tuberosity | | Ulna fac. TD | Pyram. fac. TD |
|---|---|---|---|---|---|
| | | H | APD | | |
| 57.7 | 26.4 | 40.9 | 15.1 | 18.6 | 18.9 |

**Unciform**

| TD | H | APD max. | APD min. | post. tuber. | | S.-L. fac. | | Pyram. fac. | | Mc III fac. | | Mc IV fac. | | Mc V fac. | |
|---|---|---|---|---|---|---|---|---|---|---|---|---|---|---|---|
| | | | | TD | H | TD | APD | TD | APD | TD | APD | TD | APD | TD | APD |
| 52.8 | 40.1 | 71.7 | 56.9 | 31 | 20.7 | 22.2 | 36.8 | 33.8 | 40.7 | 15.9 | 17.6 | 30.7 | 39.9 | 23.1 | 29.6 |

**Scapula**

| Tuber. Delt. DT | Diaphyse | | Ext. distal | | DT Trochl. | DAP Trochl. med. |
|---|---|---|---|---|---|---|
| | min. DT | DAP | DT | DAP | | |
| 35.2 | 30.5 | 85 | 65 | 109.7 | 65 | 86.8 |

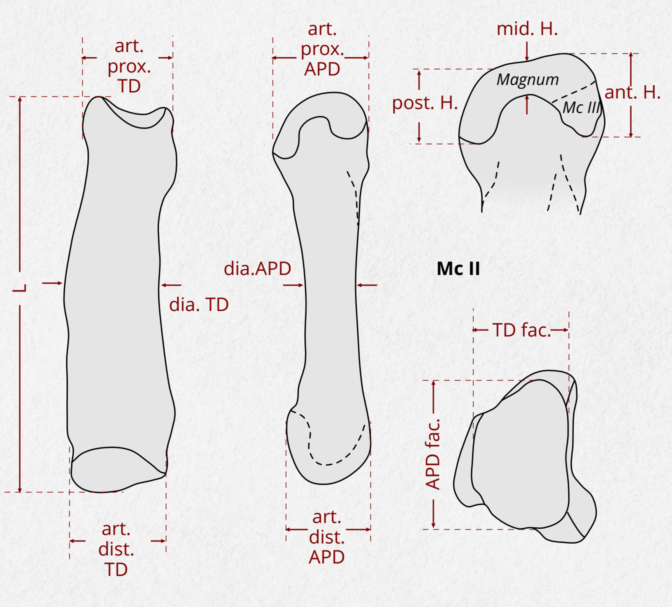


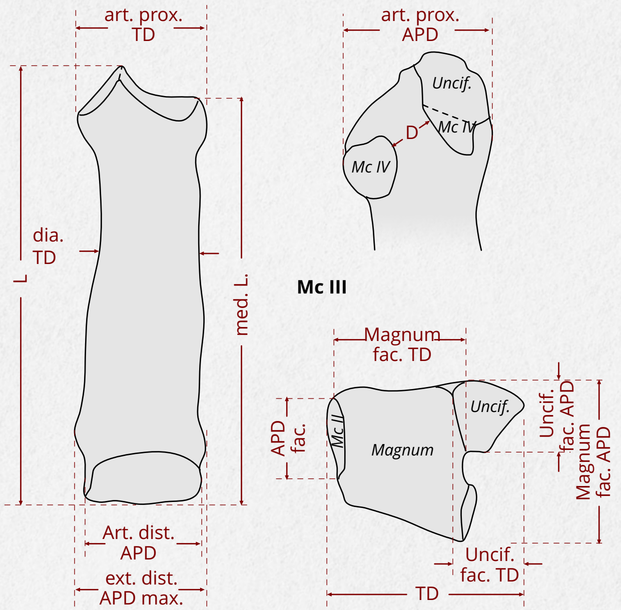


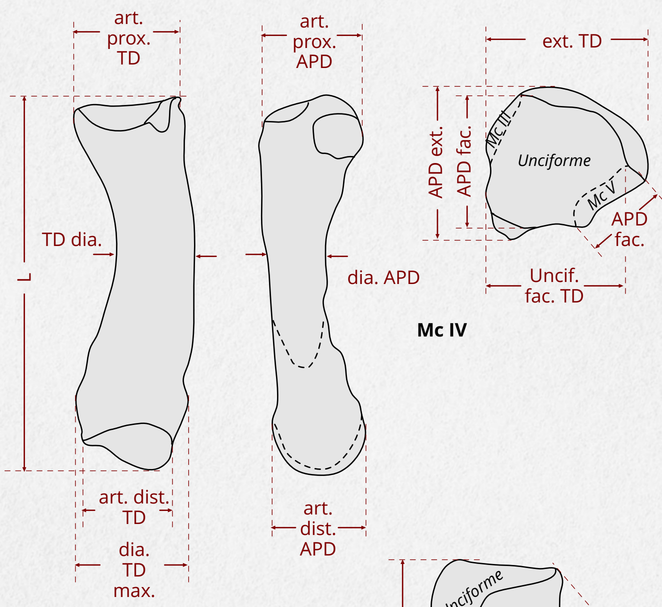


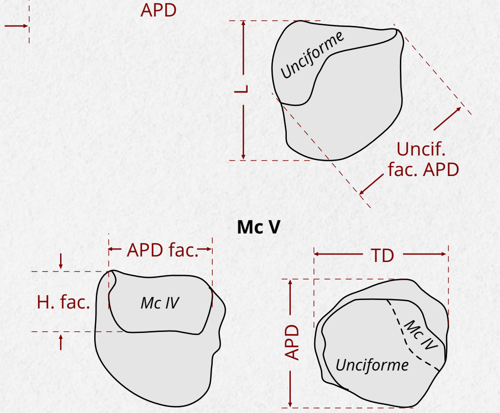


**Mc V**

| L | TD | APD | fac. Mc IV | | APD fac. Uncif. |
|---|---|---|---|---|---|
| | | | H | APD | |
| 20 | 27.2 | 25.7 | 12.9 | 19 | 22.9 |

**Mc IV**

| L | proximal art. | | fac. Unciforme | | fac. Mc V | | diaphysis (mid.) | | dist. ext. TD max. | distal art. | |
|---|---|---|---|---|---|---|---|---|---|---|---|
| | TD | APD | TD | APD | H | APD | TD | APD | | TD | APD |
| 114.7 | 38.1 | 40.2 | 29.1 | 39.6 | 12.7 | 15.3 | 32.8 | 16.8 | 43.2 | 35.1 | 36.3 |

**Mc III**

| L | L med. | proximal art. | | D fac. Mc IV | fac. Unciforme | | fac. Magnum | | fac Mc II | | diaphysis (mid.) | | dist. ext. TD | distal art. | |
|---|---|---|---|---|---|---|---|---|---|---|---|---|---|---|---|
| | | TD | APD | | TD | APD | TD | APD | H | APD | TD | APD | | TD | APD |
| 143.2 | 133.2 | 55.9 | 40.7 | 12.3 | 15.1 | 29.4 | 43.5 | 41.5 | 14.1 | 31.7 | 44 | 19.4 | 54.4 | 44.5 | 41 |

**Mc II**

| L | proximal art. | | fac. Trapezoid | | H lateral fac. | | | diaphysis | | distal art. | |
|---|---|---|---|---|---|---|---|---|---|---|---|
| | TD | APD | TD | APD | ant. | mid. | post. | TD | APD | TD | APD |
| 130.9 | 41 | 37.3 | 30.6 | 36.4 | 19.7 | 11.4 | 25.3 | 37.3 | 17.1 | 40.4 | 41.9 |

### LONG BONES OF THE HINDLIMB AND PATELLA

#### MEASUREMENTS

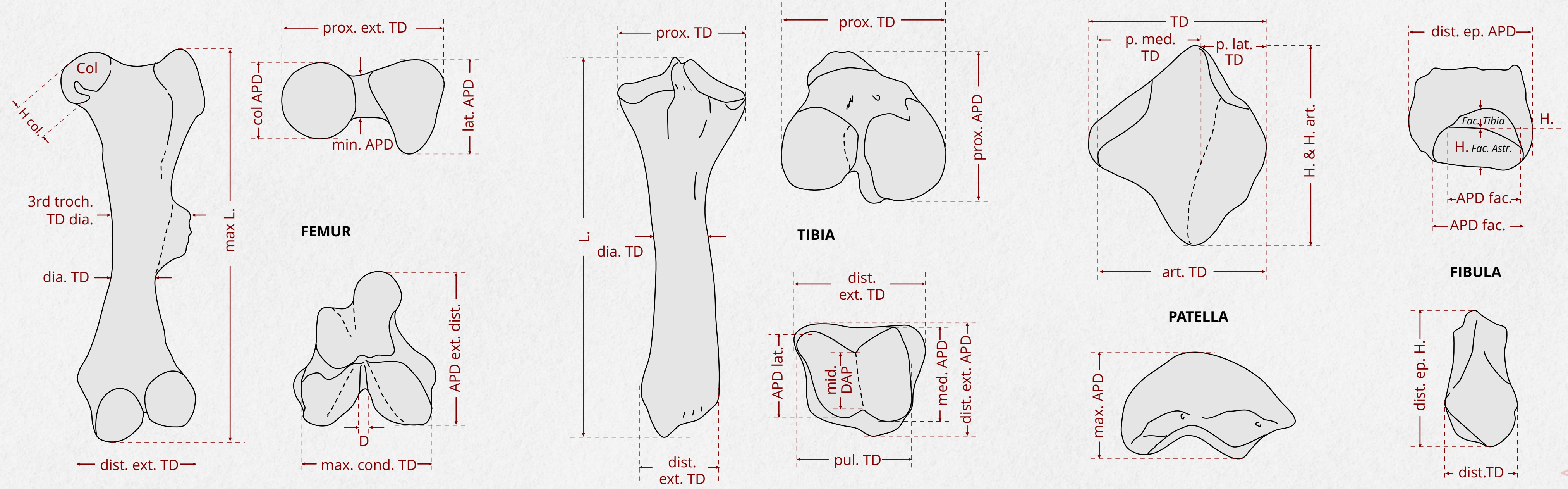


**PATELLA**

| TD | APD | | H | articulation | | TD P. med. | lateral pul. | |
|---|---|---|---|---|---|---|---|---|
| | max. | min. | | TD | H | | TD | H |
| 80.1 | 34 | 19.5 | 84 | 68.3 | 82.5 | 53.5 | 29.2 | 69.5 |

**FIBULA**

| L | proximal ext. | | fac. prox. Tibia | | diaphysis | | distal ext. | | distal epiphysis | | | fac. Tibia | | fac. Astr. | |
|---|---|---|---|---|---|---|---|---|---|---|---|---|---|---|---|
| | TD | APD | TD | H | TD | APD | TD | APD | TD | APD | H | APD | H. | APD | H |
| 267.7 | 25 | 39.3 | | | 11.8 | 14.4 | 34.2 | 41.5 | | 41.5 | | 37.1 | 9.6 | 37.1 | 18.7 |

**TIBIA**

| L | proximal ext. | | fac. med. Femur | | fac. lat. Femur | | diaphysis | | distal ext. | | Fibula fac. | | astragalus cochlea | | | |
|---|---|---|---|---|---|---|---|---|---|---|---|---|---|---|---|---|
| | TD | APD | TD | APD | TD | APD | TD | APD | TD | APD | APD | H | TD | med. APD | mid. APD | lat. APD |
| 33.3 | 115.3 | 100.5 | 57.1 | 69 | 51 | 57.4 | 47 | 40.5 | 89.5 | 60.6 | 33.6 | 9.7 | 70.6 | 50.1 | 42.4 | 48 |

**FEMUR**

| L max. | proximal ext. | | | | H col | Maj. troch. H | 3rd trochanter | | | diaphysis | | distal ext. | | condyles | |
|---|---|---|---|---|---|---|---|---|---|---|---|---|---|---|---|
| | TD | col APD | min. APD | lat. APD | | | TD dia. | TD | H | min. TD | min. APD | TD | APD | max. TD | D |
| 486.9 | 164.2 | 75.3 | 29 | 72.4 | 76.8 | 97 | 104 | 54.5 | 51.6 | 63.5 | 43 | 129.4 | 144.8 | 114.3 | 10.7 |

### THE TARSAL BONES
MEASUREMENTS

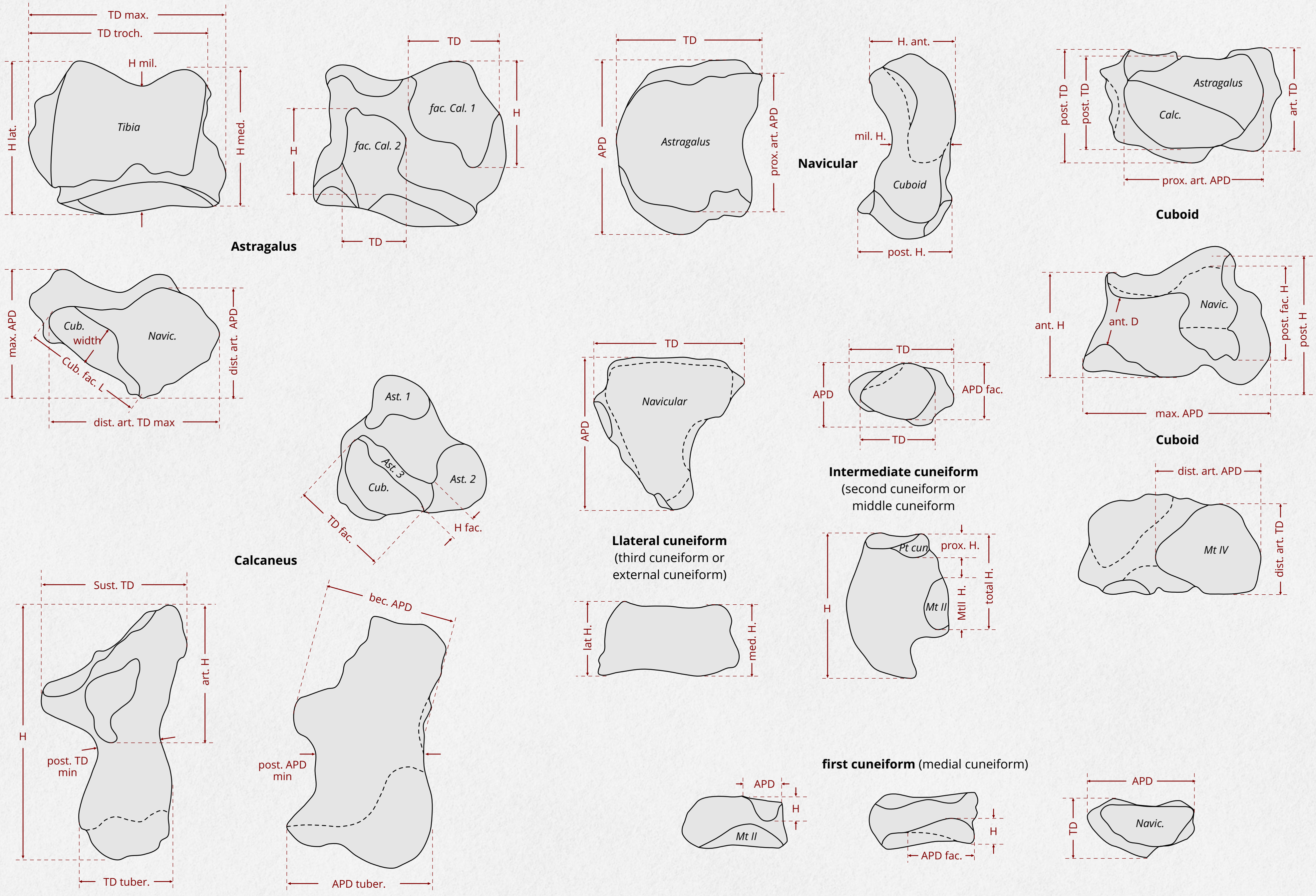

**Astragalus**

| max. TD | trochl. TD | max. APD | Height | | | Calc. fac. 1 | | Calc. fac. 2 | | distal art. | | Cuboid art. | |
|---|---|---|---|---|---|---|---|---|---|---|---|---|---|
| | | | med. | med. | lat. | TD | H | TD | H | TD max. | APD | L | width |
| 90.9 | 75.1 | 53.4 | 67.3 | 52.3 | 70.5 | 40.5 | 37.6 | 39.5 | 22.5 | 79 | 44 | 45 | 23.2 |

**Calcaneus**

| H | art. H | tuberosity | | APD bec | sust. TD | post. TD min. | post. APD min. | Astrag. fac. 3 | | Cuboid fac. | |
|---|---|---|---|---|---|---|---|---|---|---|---|
| | | TD | APD | | | | | TD | H | TD | H |
| 128.9 | 67.3 | 46.8 | 61.8 | 62.9 | 73.7 | 28.8 | 51 | 30.1 | 11.4 | 43.9 | 20.4 |

**Cuboid**

| TD | | max. APD | Height | | proximal art. | | distal art. | | medial fac. | |
|---|---|---|---|---|---|---|---|---|---|---|
| ant. | post. | | ant. | post. | TD | APD | TD | APD | D ant. | H post. |
| 43 | 31 | 61.9 | 29.9 | 49.4 | 39.6 | 43.8 | 37.3 | 37.1 | 5.8 | 26.8 |

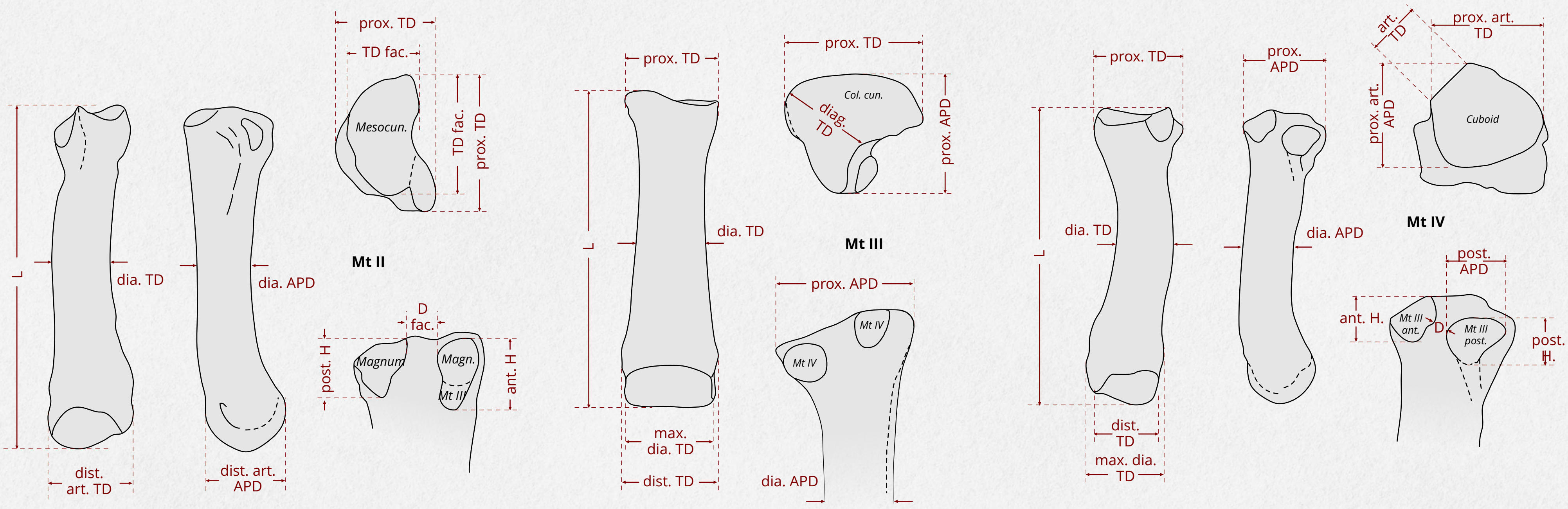


**Mt II**

| L | proximal art. | | Mesocun. fac. | | lateral fac. | | | diaphysis | | distal art. | |
|---|---|---|---|---|---|---|---|---|---|---|---|
| | TD | APD | TD | APD | ant. H. | post. H. | D | TD | APD | TD | APD |
| 106 | 31.3 | 39.9 | 25.4 | 28.4 | 5.4 | 12.9 | undivided facet | 28.7 | 19.4 | 33.8 | 39.5 |

**Mt IV**

| L | proximal art. | | | | proximal art. | | medial facets | | | | | diaphysis | | | |
|---|---|---|---|---|---|---|---|---|---|---|---|---|---|---|---|
| | TD | APD | I t | I p | TD | APD | ant. TD | ant. H | post. APD | post. H. | D | TD | APD | I t | I p |
| 101.7 | 39.1 | 42.7 | 38.4 | 41.9 | 31.7 | 33.3 | 11.3 | 16.9 | 18.7 | 14.8 | 5.5 | 30.5 | 18.1 | 30 | 17.8 |

| max. dia. TD | distal art. | | | |
|---|---|---|---|---|
| | TD | APD | TD | APD |
| 38.3 | 30.4 | 40.2 | 29.9 | 39.5 |
